# Identification of Intrinsic Features for Cortical Separability of Human and Mouse Neurons

**DOI:** 10.1101/2024.09.23.614630

**Authors:** Zhixi Yun, Wen Ye, Nan Ji, Yujin Wang, Jing Rong, Tian Li, Ruilong Wang, Yingxin Li, Hairuo Zhang, Hao Xu, Nan Mo, Binghua Song, Xin Chen, Yi Wang, Yiying Mai, Ming Ding, Song-Lin Ding, Lijuan Liu, Liwei Zhang, Hanchuan Peng

## Abstract

This work introduces a novel framework for holistic comparative analysis of cortical regions in mouse and human brains at single-neuron resolution, with a primary focus on the morphological and molecular characteristics of neurons. To do so, we generated one of the largest dendritic reconstruction datasets of cortical neurons to date, comprising 2,363 human neurons and 16,011 mouse neurons from the frontal, parietal, and temporal lobes, followed by establishing a rigorous procedure to identify anatomically and functionally corresponding brain regions with minimal variability in brain mapping. Additionally, we leveraged single nucleus/cell transcriptomic data from independent groups to validate the molecular correspondence of the brain regions identified in this study. The significance of these anatomically, functionally, and molecularly corresponding mouse-human region pairs is highlighted by examining the intrinsic features of their respective cortical regions. Our findings reveal that human neuron branching patterns differ dramatically from those in mouse brains, particularly in terms of dendritic branching frequency and normalized dendritic branching intervals. This difference is pronounced in the frontal and temporal lobes, underscoring the distinct neuronal architectures between the two species. At the single-neuron level, we found that neurons from the human frontal and parietal lobes are six times more separable than those from the same regions in mouse brains. This heightened separability is also observed between the frontal and temporal lobes, as well as between the parietal and temporal lobes in humans. We thoroughly explored the entire morphological feature space, along with its characteristic subspaces, and consistently found this distinct separability. Remarkably, this neuronal separability can be partially recapitulated when examining the global functional states of these brain lobes—using newly acquired Electroencephalography (EEG) and Magnetoencephalography (MEG) signals as physiological measures—as well as their global metabolic states, molecular profiles, and cortical geometry. These findings suggest that our comparative analysis of single-neuron intrinsic features could serve as a valuable foundation for future comprehensive studies of cross-species brain structures and functions.

## Introduction

Given the noteworthy upsurge in the number of studies that utilize mammalian neurons’ digitized morphology for brain-wide, region-specific exploration of cell typing and brain anatomy (e.g., Winnubst et al., 2019; Peng et al., 2021; Xu et al., 2021; Liu, Yun et al., 2023; Liu, Jiang et al., 2023; Han et al., 2023), a natural question arises: can the patterns primarily observed in the mouse brain be re-examined in other mammalian brains, particularly the human brain? Previous efforts have been made to compare neuronal morphology across species (Mohan et al., 2015; Benavides-Piccione et al., 2020; Mihaljević et al., 2021; Berg et al., 2021; Lee, Budzillo, et al., 2021; Bakken et al., 2021; Jorstad, Song, et al., 2023; Lee, Dalley, et al., 2023; Degl’Innocenti et al., 2023). However, three major challenges have persisted: the scarcity of reliable 3D-reconstructed neuron morphologies, limitations in brain mapping and neuron standardization techniques necessary for objective comparisons, and the lack of appropriate neuronal features to quantify differences across species. In this paper, we introduce an extensible framework to address these challenges by analyzing the morphological features and anatomical implications of single-neuron reconstructions from both mouse and human brains.

Digital atlases of mammalian brains, particularly human brains, have been recognized as essential resources in neuroscience over the past three decades (Swanson, 1995; Toga et al., 2006; Jones et al., 2009; Evans et al., 2012; Shen et al., 2012; Ding et al., 2016; Klein et al., 2017; Wang et al., 2020; Zhang et al., 2021; Qu et al., 2022; Hawrylycz et al., 2023). At the single-neuron level, these atlases are invaluable for comparing the overall distribution, projection patterns, and feature-based categorization of individual neurons across multiple spatial levels and organizational frameworks (e.g., Peng et al., 2021; Liu, Jiang et al., 2023). However, numerous challenges remain in performing cross-species comparisons, particularly between mouse and human brains. Most macroscale brain atlases are constructed primarily using Magnetic Resonance Imaging (MRI) data at millimeter or sub-millimeter resolution. For example, the Human Connectome Project (Van Essen et al., 2013; Elam et al., 2021) has mapped both the structural and functional connectivity of the human brain. In contrast, mesoscale brain atlases, such as the Allen Mouse Brain Common Coordinate Framework (CCFv3; Wang et al., 2020) and the Allen projectome of mouse neuron populations (Oh et al., 2014), provide significantly higher spatial resolution than macroscale atlases. However, there is currently no direct mapping atlas that enables single-neuron resolution comparisons between mouse and human brains.

The key idea of this work is that corresponding brain regions in mice and humans can be identified by focusing on functionally and molecularly similar areas with minimal variability in brain mapping, as validated by expert annotation. Using this approach, we compiled large single-neuron datasets from both human and mouse brains. These datasets enable us to identify corresponding regions across species and conduct a side-by-side comparison of the morphological features of neurons systematically produced in both species.

This manuscript introduces our approach to address the challenge of cross-species comparison of individual neurons in a coordinated manner. The significance of these anatomically, functionally, and molecularly corresponding mouse-human region pairs is highlighted by examining the intrinsic features of their respective cortical regions. Our findings reveal that human neuron branching patterns differ dramatically from those in mouse brains, particularly in terms of dendritic branching frequency and normalized dendritic branching intervals. We thoroughly explored the entire morphological feature space, along with its characteristic subspaces, and consistently found this distinct separability. We also found that this neuronal separability can be partially recapitulated when examining the global functional states of these brain lobes, as well as their global metabolic states, molecular profiles, and cortical folding/geometry. Our findings suggest that our comparative analysis of single-neuron intrinsic features could serve as a valuable foundation for future comprehensive studies of cross-species brain structures and functions. Our approach is generalizable using a feature analytics method, indicating its potential for a broad range of applications.

## Results

### A framework for joint mouse-human cortical analysis of single neurons in corresponding mouse and human brain regions

We used 100 whole mouse brains to generate dendritic reconstructions of 16,011 cortical neurons (**Fig. 1A**; **Methods**). We also used *ex vivo* surgical tissues of 23 human patients (**Extended Data Table 1**) to generate dendritic reconstructions of 2,363 cortical neurons of three cortical lobes (frontal lobe, temporal lobe, and parietal lobe) (**Fig. 1A**; **Methods**). To achieve these results, we leveraged our recent neuro-morphometry and neuroinformatics platforms (Liu, Jiang, et al., 2023; Han et al., 2023) for the routine generation of high-quality neuron reconstructions in both mouse and human brains. The mouse brains, which contained sparsely labeled cortical neurons, were imaged using the fMOST method (Gong et al., 2013). We then employed the All-Path-Pruning-2 (APP2) algorithm (Xiao and Peng, 2013), an automatic neuron-tracing technique, to generate initial reconstructions of the 3-D neuron morphologies from these fMOST images. These neuron models were subsequently curated using a Collaborative Augmented Reconstruction (CAR) system based on crowd-sourcing (Zhang et al., 2024). Meanwhile, for the human samples, which were obtained from patients undergoing surgeries to remove brain tumors, we utilized the Adaptive Cell Tomography (ACTomography) method (Han et al., 2023). ACTomography enabled us to produce 200 μm thick brain sections from human *ex vivo* tissues, followed by injecting Lucifer yellow into the cortical neurons and acquiring both 2-photon (2p) imaging data of local dendrites and MRI data of whole brains. Importantly, we ensured that there was no selection bias in the cortical areas studied in mouse brains by evenly sampling neurons across all regions (**Fig. 1B** – upper panel). For the human tissues, we focused on large, representative areas in the frontal, temporal, and parietal lobes corresponding to various functional regions (**Fig. 1B** – lower panel). These human neurons are mostly large pyramidal neurons from cortical layers 3 and 5, consistent with the previous study (Han et al., 2023).

With all neurons reconstructed, we next built a framework for the joint mouse-human neuron analysis. We cautioned that establishing a cross-species comparison of neuron morphologies requires careful attention to avoid overfitting the geometry during the alignment of brain anatomy. Specifically, since mouse brains lack the complex sulci and gyri found in the human cortex, forcing an alignment of these features onto a mouse brain template would be meaningless. Therefore, we initially aligned mouse brain images to a mouse brain template, and similarly for human brains. This was followed by a comparison of the neuron morphologies that had been mapped within potential corresponding regions of their respective canonical spaces. Concretely, we employed mBrainAligner (Qu, et al., 2022), a landmark-based cross-modality registration tool, to map all individual brains to their respective template brains (**Fig. 1A** – step S1). For mouse brain images, we used the CCFv3 as the template. For human brains, we utilized the ICBM2009c nonlinear template (Fonov et al., 2009) which led to enhanced alignments compared to previous alignments (Han et al., 2023) based on the MNI152 linear template. Subsequently, mBrainAligner was used again to map individual neuron reconstructions or *ex vivo* tissue segmentations to the two respective atlases (**Fig. 1A** – step S2; **Methods**). Given that the human *ex vivo* tissues were extracted from a limited number of cortical regions (**Fig. 1B**), we identified the corresponding mouse brain regions for these human samples based on expert neuroanatomists’ annotations of brain regions corresponding to similar brain anatomies and functions, as well as based on a data-driven approach using gene expression analysis developed below (**Fig. 1A** – step S3; Methods). Within the canonical spaces of these atlases, standardized neuron morphologies could be quantitatively compared based on various features (**Fig. 1A** – step S4).

We examined the global morphological features of the standardized mouse and human neurons to ensure they could be compared fairly and quantitatively. We found that the dendritic morphologies of mouse neurons in this study statistically matched those produced in an earlier study (Peng, et al., 2021) in terms of their global morphometry (**Fig. 1C**). Our human neurons exhibit less overall reconstructed neurite length than mouse neurons (**Fig. 1C; Extended Data Fig. 1A**), attributable mostly to the common situation of truncated neurites in brain tissue sections (Han, et al., 2023). The detailed comparison of neuron morphology will be discussed later in the paper.

**Fig. 1.**
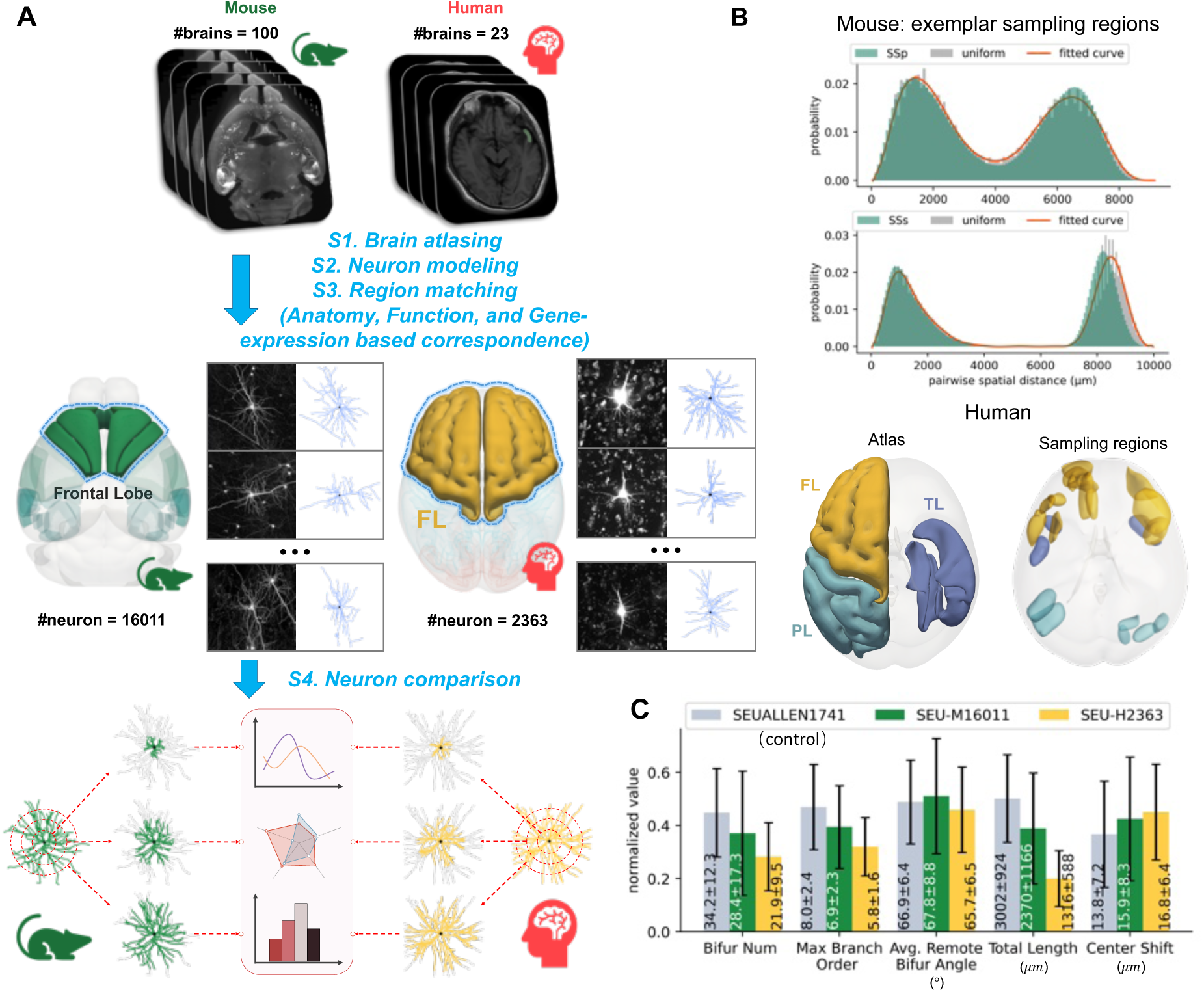
A joint atlas of mouse-human cortical neurons along with dendritic morphometry. **A**, The four major steps, S1 ∼ S4, in assembling the joint atlas. **B,** Illustration of the cortical regions from which neurons were reconstructed. For the mouse brains, the pairwise spatial distance distribution shows consistency across reconstructed neuronal somas and uniformly sampled points in cortical regions. **C,** Comparison of morphological features of the mouse (SEU-M16011) and human (SEU-H2363) neurons, those from a control dataset of mouse neurons (SEUALLEN1741, as in Peng et al., 2021) within a neighboring sphere (radius = 100μm) from the respective somas. The bar plot shows the mean values of morphological features and their respective standard deviations.

### *De novo* identification of statistically corresponding cortical regions matches anatomical annotation of brain lobes with similar functions

For an effective comparison of neuron morphology across species, we generated several references as baselines to determine the significance of the comparison. To address the intrinsic anatomical differences between mouse and human brains, we registered the collected brain images of both species to their respective atlas templates. We then computed the anatomical average and variation maps for each species, which helped us identify brain areas with sufficient registration precision for further single-neuron morphometry and statistical analysis (**Figs. 2 and 3**).

For SEU-M16011, we registered the respective fMOST images to CCFv3 (**Methods**). The overall average map of the registration was anatomically consistent with the mouse CCFv3 atlas. Indeed, the variance is less than 0.05 in most brain regions, especially the cortical regions studied here (**Fig. 2A**). Indeed, although the mouse neurons and their neurites span the entire cortical regions of the frontal, temporal, and parietal lobes (FL, TL, and PL), their somas and neurite arbors exhibit the greatest density in the middle of these lobes. This distribution is not influenced by the slightly higher registration variation found around the borders of some brain areas. (**Fig. 2A**). Out of the 16 mouse cortical regions in the three lobes, supplemental somatosensory area (SSs), visceral area (VISC), and postrhinal area (PoR) are the three brain regions that contain the largest overall density of mouse neurons and demonstrate the greatest arbor density (**Fig. 2B**). The normalized variation of registration for all 16 brain regions is around 0.02, indicating that the morphometry of cortical neurons in SEU-M16011 is less likely to be impacted by potential registration inaccuracies and anatomical variability of original brain images (**Fig. 2C**).

For SEU-H2363, we mapped the MRI data of all 23 patients to the ICBM2009c template (Fonov et al., 2009) (**Methods**). Before brain-image registration, surgeons annotated the names of the *ex vivo* brain regions and also labeled their actual 3-D segmentations in the sample extraction areas. After the registration, these standardized locations of brain regions reveal the overall distribution of sampled neurons in the human atlas (**Fig. 3A**). We found that the surgeon-generated annotations of brain regions matched with annotations based on the CerebrA atlas (Manera et al., 2020) (**Fig. 3A, 3E**), which was a more accurate version of Mindboggle-101 atlas (Klein and Tourville, 2012). The registered cortical regions of interest also display overall small anatomical variations of less than 0.06 (**Fig. 3B**). Indeed, the intensity variances of the cortical regions all peak at close to 0.02 (**Fig. 3C**), comparable to those of the registered mouse brains (**Fig. 2C**). For all three cortical lobes, we also sampled neurons at a rate of approximately 100 neurons per subject per lobe (TL: 104.8, FL: 94.9, and PL: 116.7) (**Fig. 3D**). This implies that the neurons randomly sampled from these regions are not expected to display a systematic skewness attributable to brain registration or sampling ratios in subsequent analyses. Finally, the sampled human neurons show clear dendritic arborization, making them suitable for cross-species comparison. As we employed expert annotators to reconstruct the dendritic arbors as completely as possible, subject to the visibility of neurite signals in the 2-photon images of dye-injected cells, the 3-D reconstructed dendritic arbors can be as long as almost 9mm (**Fig. 3A**). Although human neuron reconstructions were not registered into a canonical space, comparisons with registered and non-registered mouse neurons on morphological features could be used to speculate that the morphological differences are likely very small in human neurons (**Extended Data Fig. 2**). Considering the neurite lengths of human neurons could be under-represented, the key dendritic branching patterns, such as fully traced branches and critical furcation points, remained suitable for the analysis of key neuro-morphological patterns in this study.

To utilize the data more effectively and achieve a deeper analysis, we determined corresponding regions based on the anatomical and functional correspondences between mouse and human brain regions. Initially, we identified the lobes to which human brain regions belonged and matched these lobes to anatomically and functionally corresponding regions in the mouse brain. We then further refined this correspondence by matching human and mouse brain regions at a finer level according to well-established anatomical or functional knowledge, which allowed for more detailed and targeted cross-species comparisons of neuronal morphology (**Fig. 3E, Extended Data Table 2**).

**Fig. 2.**
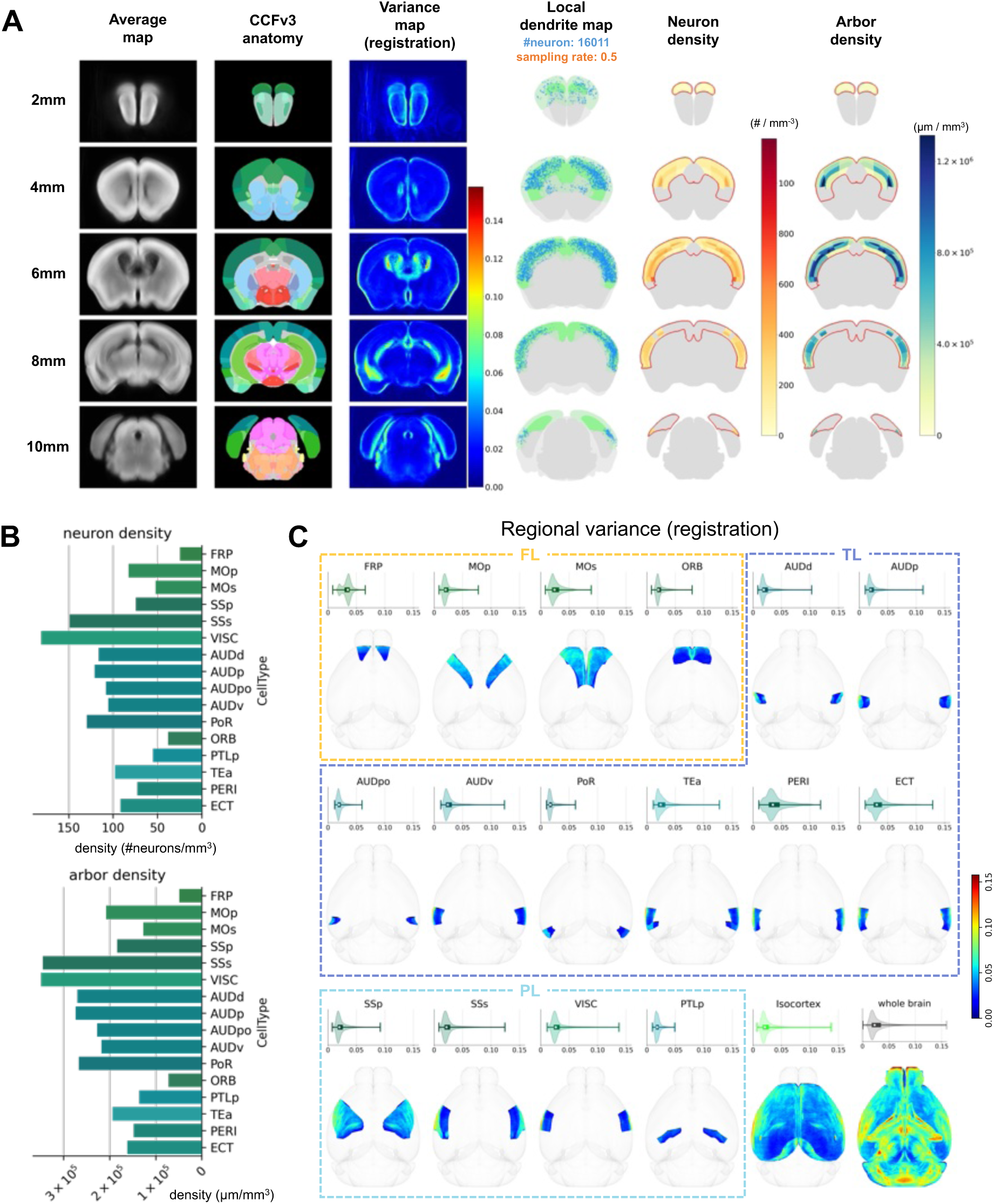
An atlas of mouse brains: anatomical average and variance maps produced in registration, along with the density patterns of reconstructed neuron-dendrites (SEU-M16011) registered onto the CCFv3 template. **A**, Six uniformly-spaced coronal image series along the anterior to the posterior ranging from 2mm to 10mm in the CCFv3 space. First panel: average grayscale map of 100 registered brain images. Second panel: the reference CCFv3 anatomical atlas colored by brain regions. Third panel: intensity variance of 100 registered brain images. Fourth panel: local arbor reconstructions of cortex mapped onto the CCFv3 atlas; A random sample of 50% of the neurons was used for whole-brain visualization. Fifth panel: neuronal density of reconstructed neurons in cortical regions. Sixth panel: cortical arbor density quantified by branch lengths per unit volume. The cortical regions were outlined in red in the fifth and sixth panels. **B,** Neuron density and arbor density in regions of interest. **C,** Registration variance of specific cortical regions, cortex, and whole brain. The distribution of normalized intensity variance for each brain region was shown in the violin plot. In addition, regional variance was displayed along the dorsal-ventral axis using maximum intensity projection. Brain regions were framed into different human lobes based on their function.

**Fig. 3.**
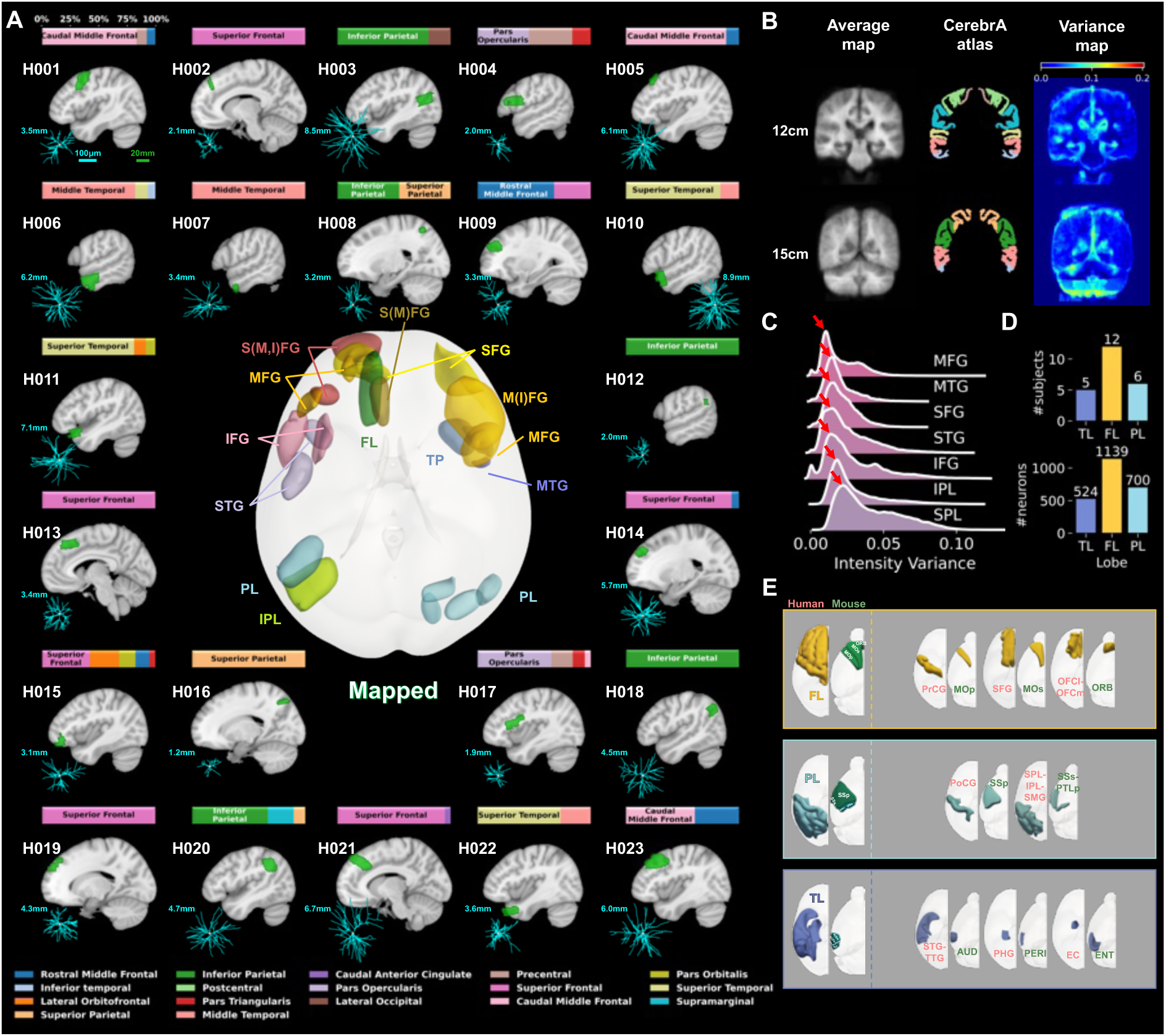
An atlas of human cortical neurons, where anatomical average and variance maps were produced in registration (target = ICBM 2009c Nonlinear Asymmetric template), along with examples of reconstructed neuron-dendrites (SEU-H2363). **A**, Visualization of 23 sampling regions on the sagittal slice of ICBM2009c template. The sampling region was annotated in green. The bar graph at the top of the brain image showed the volume proportion of each brain region in the sampling region. The value of the morphological feature “Total length” was labeled beside the neuron example. Scale bar in cyan for neuron reconstructions, 100μm. Scale bar in green for brain images, 20mm. **B,** Overall registration results of 23 human brain images with the CerebrA anatomical atlas in coronal view at 12 mm and 15 mm positions along the anterior-posterior direction. First panel: average grayscale map of registered brain images. Second panel: the reference CerebrA atlas colored by brain regions. Third panel: variance of normalized voxel intensity (range: 0∼1) of registered brain images. **C,** The variance distribution of normalized voxel intensity in each representative region. The peaks were marked with red arrows. **D,** Number of subjects and number of neurons encapsulated in each lobe. **E,** Expert-anatomist annotation of brain regions that share corresponding functions in human and mouse brains. See **Methods** for full names of brain region abbreviations.

### Cellular RNA sequencing validates key corresponding mouse-human regions in frontal, parietal, and temporal lobes

While comparing anatomical and functional properties provided insights into the correspondence of cross-species brain regions, we further explored genetic correspondence by utilizing transcriptomic data. Despite potential differences in gene expression profiles, most cell types were conserved across species. By comparing the cellular composition and proportions of shared cell types in the human and mouse brain regions, we constructed a cellular-composition similarity map and inter-regional genetic similarity rankings to assess and validate regional correspondence further.

We considered two state-of-the-art whole-brain transcriptomic datasets: single-cell RNA sequencing (scRNA-seq) data from mice (Yao et al., 2023) and single-nucleus RNA sequencing (snRNA-seq) data from humans (Siletti et al., 2023). The raw gene expression matrices were filtered to exclude non-neuronal and non-cortical cells, and then 15,506 homologous genes (**Extended Data Table 3**) common to both species were queried from the National Center for Biotechnology Information (NCBI) Gene database (National Center for Biotechnology Information, 2004). After quality control, we integrated these datasets based on the preprocessed gene expression matrices to eliminate the batch effects (**Fig. 4A; Methods**). The cells were well-mixed across species (**Fig. 4A**) and across shared well-annotated cell types reported in the original datasets (**Fig. 4B, left panel**). This integration allowed for further unsupervised clustering (**Fig. 4B, right panel**) and cell counting in each cluster. The gene profiles of each brain region were able to be encoded in proportions of clusters which indicated various cell types (**Methods**).

Using the normalized cell-type proportions of each region, we constructed a correlation map to approximate pairwise genetic correspondence (**Fig. 4C; Methods**). Due to the potentially distinct proportions of cortical cell-types in mice and humans (Bakken et al., 2021; Fang et al., 2022; Chartrand et al., 2023), normalization of cell-type proportions was performed by within-species (**Methods**), which assumed that the relative changes of the same cell type were consistent between different brain regions within a species. To improve robustness, the correlation map also provided a confidence value to indicate the stability of the correlation scores computed with different numbers of principal components (**Fig. 4C; Methods**). In this map, region pairs with high correlation and high confidence indicate possible genetic correspondence. For instance, the middle frontal gyrus (MFG) and inferior frontal gyrus (IFG) in the human brain correlate well with the somatomotor areas and frontal pole (MO-FRP) in the mouse brain (mean ± s.d.: 0.47 ± 0.04, 0.54 ± 0.02). The superior parietal lobule (SPL), visual areas and posterior parietal association areas (VIS-PTLp) also match well (mean ± s.d.: 0.59 ± 0.02). The cuneus-lingual (Cun-LiG) and visual areas (VIS) show a strong correlation (mean ± s.d.: 0.73 ± 0.01), which also indicates that mice and humans might be more conserved for vision-related cellular mechanisms. However, due to the difficulty of microdissecting small mouse regions for scRNA-seq, some cells from neighboring regions were mixed, making cell-type proportions in these regions unrepresentative of individual brain regions and potentially introducing imprecise genetic similarities.

To provide a more robust delineation of genetic correspondence, we adopted three metrics and used an integrated rank instead of individual metric values to avoid situations where the values of multiple metrics cannot be simply averaged (**Methods; Extended Data Fig. 4**). Relevant regional correspondences (**Extended Data Table 2**) were thoroughly examined based on the available transcriptomic data. The anterior cingulate area (ACA) in the mouse brain best matched the rostral anterior cingulate cortex in human brain regions. The VIS in the mouse brain best matched the Cun-LiG or Cun in human brain regions (**Fig. 4D**). We also note that the differences in neuron composition between VIS sub-regions (e.g., V1 and V2) could be significant (Bernard et al., 2012; Jorstad, Close, et al., 2023). It will be interesting to explore these nuances further as higher spatial resolution data become available. Other anatomical and functional corresponding regions also showed the highest or very higher ranks except for the pair of the primary motor area (MOp) and precentral gyrus (PrCG) which might have been affected by non-identical batch samples (Siletti et al., 2023; **Extended Data Fig. 4**). These approaches enabled us to determine the relative similarity of cell type proportions across all compared regions, strengthening the anatomical and functional correspondences and making them more convincing.

**Fig. 4.**
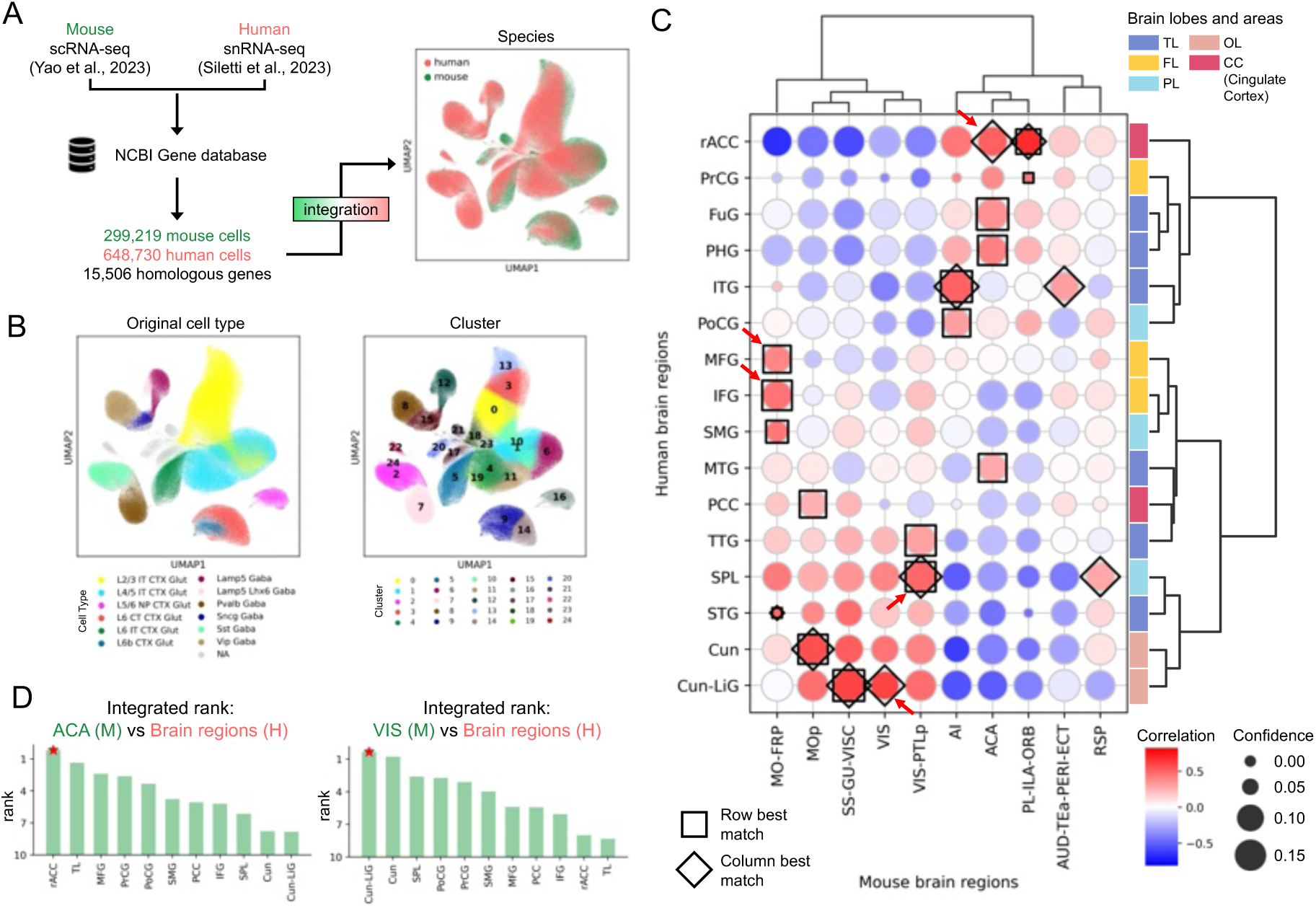
Measurement of cross-species genetic correspondence based on cell-type composition. **A,** Homologous gene determination and integration of mouse and human transcriptomic data. UMAP graph shows the distributions of cells colored by species. **B,** UMAP results on integrated data labeled by shared cell types in both datasets and their clustering results. Abbreviations: L: layer; IT: intratelencephalic; CTX: cerebral cortex; Glut: glutamatergic neuron; CT: cortical-thalamic; NP: near-projecting; Gaba: GABAergic neuron; NA: not applicable, includes the rest cell types. **C,** Clustering map illustrating genetic similarity between human and mouse regions. Point colors represent scores derived from averaging Pearson correlations for various combinations of cell-type compositions between human and mouse regions. Point sizes indicate the confidence of the correlation value. Square shapes denote the maximum values in each row, while diamond shapes indicate the maximum values in each column. The right sidebar shows lobes and brain areas to which human brain regions belong. **D,** Bar plots showing integrated ranks of similarity in cell-type compositions between specific mouse and human brain regions under multiple similarity metrics. Rankings are sorted in descending order. Red stars indicate potential functionally corresponding brain regions in humans relative to specific mouse brain regions. See **Methods** for full names of brain region abbreviations.

### Human and mouse cortical neurons differ in branching frequency and intervals especially in the secondary motor cortex

To make the most of the limited data and explore the morphological similarities and differences between human and mouse cortical neurons, even at finer regional correspondences, we developed an analytical framework that includes multi-level morphological characterization and soft-labeled neuron analysis based on robust regional correspondences.

We initiated our morphological analysis on the species level to gain a holistic understanding of neuronal characterizations. Typically, our knowledge of complex structures begins with the examination of the basic constituent structures. Therefore, we first investigated the characteristics of single branches. In consideration of the completeness of neuron reconstructions, we focused on the non-terminal branching patterns of lower branch-order (BO) branches, ensuring these were fully traced from limited image signals. We found that, when pooling single branches from all neurons, the peaks in the branch length distributions differed minimally between the two species in low BO branches (BO2, BO3, BO4; **Fig. 5A upper panel**). This might indicate similarity in signal transmission between human and mouse at lower-order single branches. However, it was known that soma volume correlates with dendrite volume in the mouse cortex (Gao et al., 2023). In addition, the total length of basal dendrites in human temporal lobe neurons was approximately twice that of mice (Mohan et al., 2015). Considering soma sizes of human neurons were on average 1.66 times larger than those of mouse neurons in our study (**Extended Data** Fig. 1C), we normalized mouse neurons by scaling them using the average soma radius ratio between species in order to eliminate the potential effect of overall dendrite size on comparison between the two species. Under these circumstances, the distributions of branching lengths are much more distinguishable (**Fig. 5A upper panel**). In other words, considering the overall scale in the human brain, neurons tend to use relatively short, low-order branches to bifurcate rapidly. We also examined sibling branches, defined as pairs of branches connected to the same parent branch, by calculating the angle between them. The peaks in the two angle distributions are nearly identical (**Fig. 5A upper panel**). Neither of these features at the level of the single-level branch shows obvious differences between lobes within the species. (**Extended Data Fig. 5**).

Building on our analysis of the single-branch structures, we further examined the overall structure of local neurons. Minor variations are observed between species when focusing on single or sibling branches. However, noteworthy differences emerge when examining the “bifurcation numbers” and “total length” within a local spherical range near the soma (radius = 50μm, with an “R50_” prefix before the morphological feature name). This range is moderate for encompassing adequate branches while minimizing the inclusion of incomplete branches resulting from insufficient slice thickness (**Methods; Extended Data Fig. 1A**). Human neurons have a higher number of branches compared to mice, even though human neurons theoretically contain fewer branches due to their larger cell body size within the same spherical range. (**Fig. 5A lower panel**). Based on the same normalization approach (**Methods**), a more intuitive discrepancy in bifurcation numbers and total length between the two species emerge (**Fig. 5A lower panel**). Further examination of branch numbers of the first five BO indicates that the branch abundance of human neurons is mainly due to richer primary branches (BO1). This analysis suggests that while the basic structure of low-order branches is conserved between species, the way these structures are assembled highlights the key differences. In particular, human cortical neurons exhibit a more complex initial branching pattern compared to their counterparts.

To further understand morphological relationships at a finer anatomical level, we conducted a more targeted comparison between the cross-species corresponding regions (**Fig. 3E**). Given that some human brain sampling regions spanned multiple areas, complicating accurate annotation by the expert-anatomist, we used a probabilistic region assignment method to allocate soft labels to neurons based on the volume proportion of sampled tissue in different brain regions (**Fig. 3; Extended Data Fig. 3A**). Soft labels maintain consistency with manual labels and also provide detailed information on regional localization.

Combining probabilistic labels with our corresponding region analysis, we calculated the weighted mean and cumulative probability quartiles of neuronal morphological features in each region (**Fig. 5C; Methods**). To ensure meaningful feature representation, we selected regions with sufficient neuron numbers and adequate cumulative probabilities (**Methods; Extended Data Fig. 3B**). We examined bifurcation numbers, the first five BO branch numbers, and max branch order across corresponding regions. Notably, secondary motor area (MOs) and superior frontal gyrus (SFG) exhibit great discrepancies in branch abundance, with human SFG neurons displaying richer and more complex branching structures. This difference is likewise due to more first-order branches, indicating a more complex signal integration, possibly underscoring advanced executive functions and higher-order motor planning capabilities in humans. Notably, this approach reduces the potential inaccuracies and rigidity of manual region labeling, enabling more precise and reliable comparisons of neuronal morphologies.

Our data, as discussed above, indicate that the conventional metrics for comparing human and mouse neurons is overly simplistic, failing to capture the variation in normalized dendritic branching intervals. Additionally, the frequency of dendritic branching within a comparable volume differs considerably between these two species. Both crucial features have not been rigorously quantified in previous studies.

**Fig. 5.**
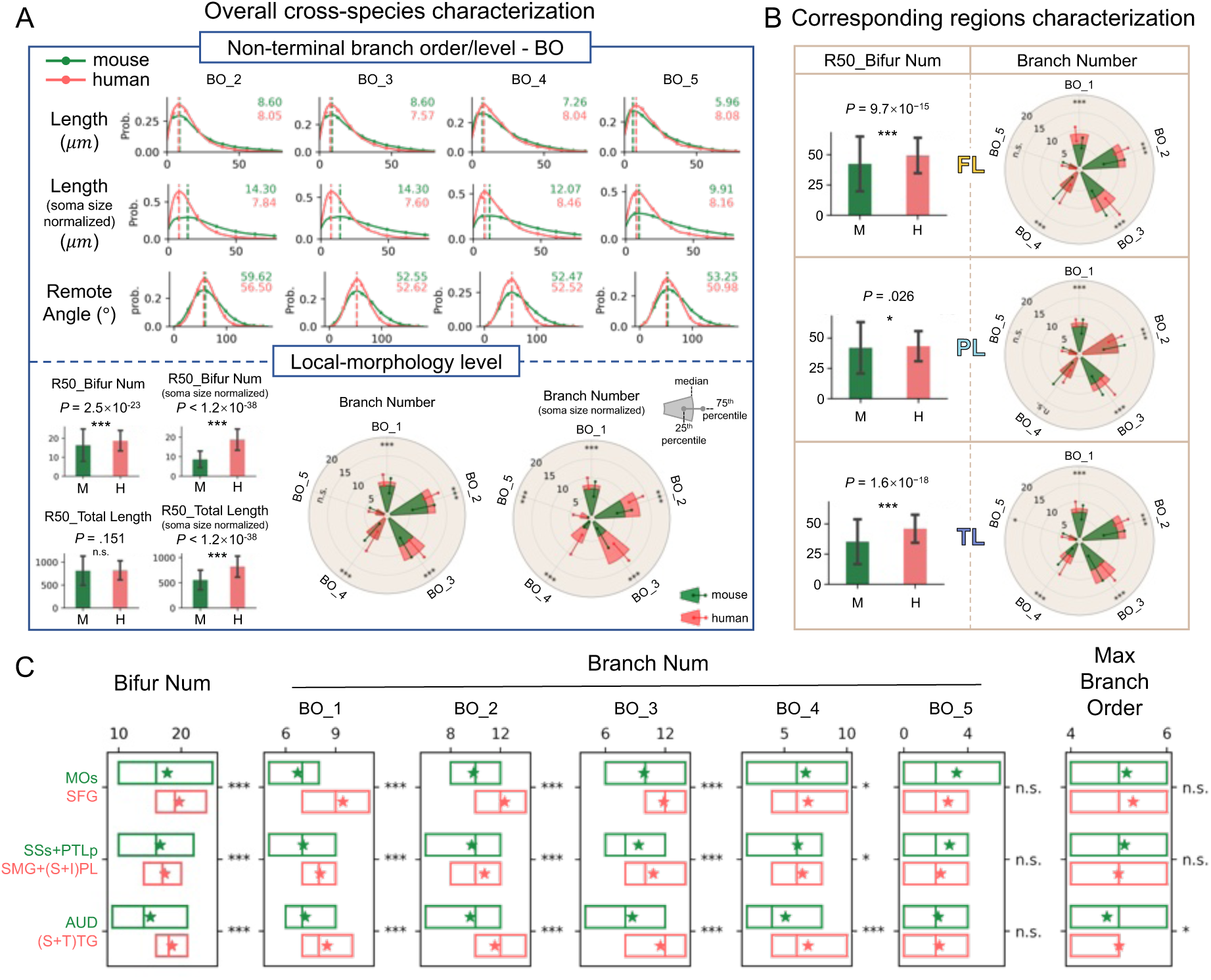
Cross-species morphometric analysis through multigranular correspondence. **A**, Comparison of morphological features of human and mouse cortical neurons. Upper panel: Distribution of single-branch and sibling-branch morphological features pooled by branch orders at the level of non-terminal branches. The dashed line perpendicular to the x-axis represents the peak of the distribution, with actual values labeled in the upper right corner (mouse in green and human in red). Lower panel: Morphological features of local neurons. Bar plots display average values with error bars showing standard deviation. P values and significance levels are labeled (two-sided Mann-Whitney U test, *P<.05, ***P<.0001; n.s.: not significant, P≥.05). Radar plots depict the median value (edge of the sector) and the 25th to 75th percentiles (points on either side of the line). Significance levels are labeled (two-sided Mann-Whitney U test, *P<.05, ***P<.0001; n.s.: not significant, P≥.05). **B,** Comparison of morphological features of cortical neurons in human and mouse anatomical corresponding lobes. **C,** Comparison of morphological features of cortical neurons in human and mouse functional corresponding regions. Box plots illustrate the distribution of morphological features, with the central line representing the weighted median, and the box extending from the first to the third weighted quartile. The star denotes the weighted average. Significance levels are labeled (two-sided weighted Kolmogorov–Smirnov test, *P<.05, ***P<.0001; n.s.: not significant, P≥.05; **Methods**). Complete P values are recorded in **Extended Data Table 4**. See **Methods** for full names of brain region abbreviations.

### Human dendrites are far more distinct across major cortical lobes compared to those in mice

Neurons from the cortical lobes of both human and mouse brains show remarkable differences in local branching morphometrics (**Fig. 5**). However, the distinctions between neurons from different lobes within the same species remain unclear. To explore this, we used a quantitative approach, measuring morphological discrepancies of neurons across different lobes within each species using classification accuracy (**Methods**). We reduced data variability and minimized the risk of overfitting by using five-fold cross-validation and optimized classifier parameters through grid search, resulting in balanced accuracy scores and other categorical evaluation metrics (**Methods; Extended Data Fig. 6**).

As more features are incorporated into the training, the local morphologies of the three human lobes (FL, PL, TL) consistently show stronger discriminatory power between each pairwise combination compared to those in mice (**Fig. 6A, 6B, 6C, upper panels**). For the “FL-PL” pair, human neurons achieve approximately 70% classification accuracy, while mouse neurons reach only around 55% (**Fig. 6A, upper panel**). We used multivariate Jensen–Shannon divergence (MJSD) to quantify separability (**Methods**). The difference of this separability could reach approximately 6.18 times (**Extended Data Table 5**). Although the scores for “FL-TL” and “PL-TL” are lower, human neurons still exhibit higher classification scores than mice. This result suggests that the spatial organization and structural complexity of local morphologies vary more significantly across human cortical lobes compared to those in mice.

Higher classification performance generally indicates more separable features. We thus analyzed the normalized distribution discrepancy (NDD) of features between neurons from every two lobes successively (**Fig. 6A, 6B, 6C, middle panels**). We counted the number of features where the discrepancy in humans was greater than that in mice. For the “FL-PL” pair, the NDDs of all 21 morphological features in human lobes are higher than in mouse lobes, indicating greater differences between FL and PL in humans for nearly every morphological characteristic. In the other two pairs of lobes, the NDD scores for more than half of the human features are higher than the NDD scores for the corresponding features in mice.

To facilitate a fair comparison, we also investigated the morphological separability using representative feature subsets for both species. With the minimal-redundancy-maximal-relevance feature selection algorithm (mRMR, Peng, et al., 2005), we identified the top five most characterizing features for each lobe pair in both species. In most cases, humans have higher NDD scores for these top-ranked features. Even when using the top-ranked mouse features as references, few mouse features have higher NDD scores than humans. For the “FL-TL” and “PL-TL” pairs, three and two out of five top-ranking features have higher NDD scores in mice, respectively (**Fig. 6A, 6B, 6C, lower panels, scatter plots**). In the meantime, we noted that top-ranking features within an individual species rarely overlapped between species. There is only one shared feature in the “FL-TL” (Avg. straightness) and “PL-TL” (3D Density) pairs (**Fig. 6A, 6B, 6C, lower panels, texts in bold**).

**Fig. 6.**
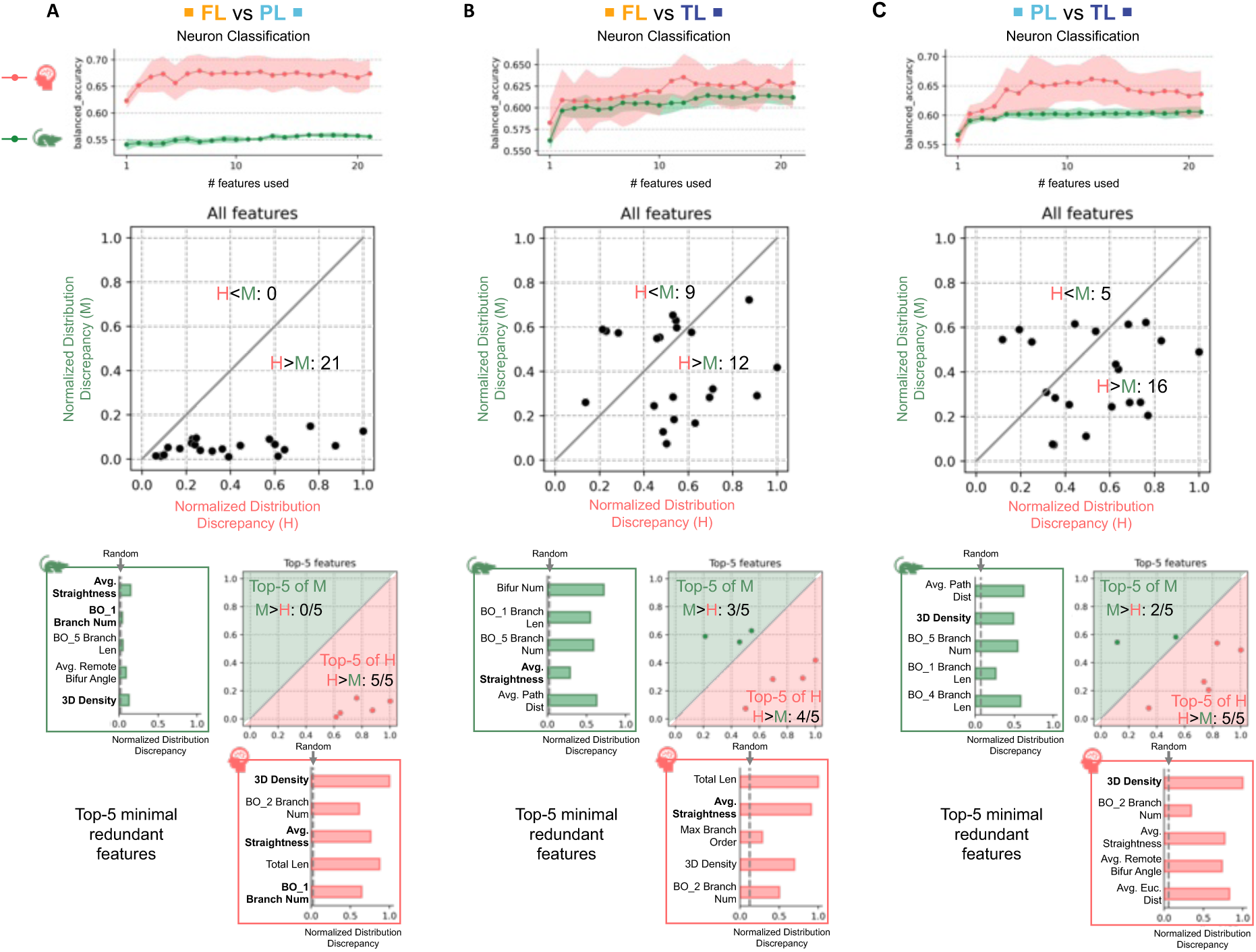
Neuronal classification accuracy and distributional differences in morphological features between lobes within species. **A**, Upper panel: Line plot showing classification accuracies of neuronal morphologies between the frontal and parietal lobes using iteratively added features. The line represents the average balanced accuracy derived from 5-fold cross-validation. The shaded area represents the mean ± standard deviation. Middle panel: Scatter plot showing the normalized distribution discrepancy (NDD) of the same morphological features between the frontal and parietal lobes in human and mouse neurons. The NDD is calculated using Jensen–Shannon divergence, with the maximum normalized to 1. The diagonal line represents y=x; points below the line indicate higher distribution discrepancies in humans, and points above the line indicate higher distribution discrepancies in mice. Lower panel: Joint graph showing the NDD of the top-5 minimal redundant features selected by the mRMR algorithm in each species. The scatter plot is as described in the middle panel, but each species is shown only in the half of the region where its characteristic differences predominate. The bar plot shows the NDD of the top-5 features, with a dashed line indicating the NDD of two random distributions as a control. Features highlighted in bold font are those appearing in the top 5 features for both species simultaneously. **B,** Comparison between the frontal and temporal lobes with the same elements as in **panel A**. **C,** Comparison between the parietal and temporal lobes with the same elements as in **panel A**.

### Human cortical lobes exhibit varying separability in multi-modal space

Given the quantitative evidence that human cortical lobes are more separable than mouse brain lobes in the neuron morphology space, an interesting next question is how these morphological features compare to other cortical distinctions measured through anatomical, physiological, or biochemical methods. To this end, we focused on human brain only and explored separability using molecular profiles (Siletti et al., 2023), cortical anatomical geometry information including cortical thickness and curvature (**Supplementary Material 4**), physiological signals such as EEG and MEG data of newly acquired resting state and visual tasks (shape-stimulation) (**Methods**), and global metabolic states (Aanerud et al., 2012; **Supplementary Material 5**). At the molecular level, we investigated gene expression patterns across cell populations that vary between lobes. For physiological signals, we examined the signal characteristics of different sensors corresponding to specific visual tasks or resting state (**Methods**).

Overall, we found that different modalities reflected varying degrees of lobe separability (**Fig. 7A; Methods**). Based on MJSD measurements, morphology-based separability of human cortical lobes is intermediate between gene expression and physiology-based metrics. Gene expression exhibits the highest degree of separability, while physiological signals are more state-or task-specific, tending to show separability between lobes only in task-dependent contexts (e.g., MEG_circle and MEG_triangle tasks) (**Fig. 7A**).

We then examined the relative separability of cortical lobes across different modalities to determine whether certain data types or features were more effective in distinguishing specific pairs of lobes. To assess the separability among all three pairs of lobes, we visualized their ternary relationship using a circle, where the length of the edge connecting two lobes reflects their relative separability—the longer the edge, the greater the distinction between them. (**Fig. 7B-D**). Importantly, we also showed the total MJSD scores at the center of the circle: a low score indicates poor overall separability between the lobes, making the relative differences less meaningful.

In this analysis, our finding is that single-neuron morphology provides a more balanced classification of cortical lobes (**Fig. 7B**). As we analyzed most pyramidal neurons’ morphologies, which are also excitatory typically, it is reasonable to hypothesize that this type of neurons plays a role in differentiating cortical lobes. Interestingly, gene expression profiles show a stronger ability to distinguish between FL and TL when all neurons are considered (**Fig. 7C-left**). When we focused solely on excitatory neurons, the distinction becomes even more pronounced (**Fig. 7C – middle**), supporting our hypothesis that differences in excitatory neurons may be a factor in distinguishing these cortical lobes. EEG data, while effective at distinguishing FL-PL pairs, show lower overall lobe separability (**Fig. 7D**). MEG data, on the other hand, exhibit varying separability depending on the task, with stronger FL-TL differentiation during the ‘circle’ task and better FL-PL separation during the ‘triangle’ task (**Fig. 7D**). For metabolism (Aanerud et al., 2012), we analyzed both cerebral blood flow (CBF) and the cerebral metabolic rate of oxygen consumption (CMRO2) (**Supplementary Material 5**). The relative magnitudes of these metabolic features are consistent across age groups (**Fig. 7E**). Their overall distributions also show little differentiation between the lobes (**Fig. 7E**). Our quantitative evidence suggests single-neuron morphology may offer more robust and generalizable classification of cortical lobes compared to task-specific physiological signals.

As a global reference, we also examined the cortical geometry in characterizing these three lobes. We used four metrics—average thickness, thickness standard deviation, integrated rectified mean curvature, and integrated rectified Gaussian curvature—to assess geometric differences. These geometric features were calculated both at the level of specific anatomical regions and at the broader lobe level (marked with a plus symbol on each box plot in **Fig. 7F**) using the ICBM2009c human brain template (**Methods; Supplementary Material 4**). Our analysis shows that the parietal lobe has a relatively smaller average thickness, while the temporal lobe displays reduced curvature compared to the other lobes (**Fig. 7F**).

**Fig. 7.**
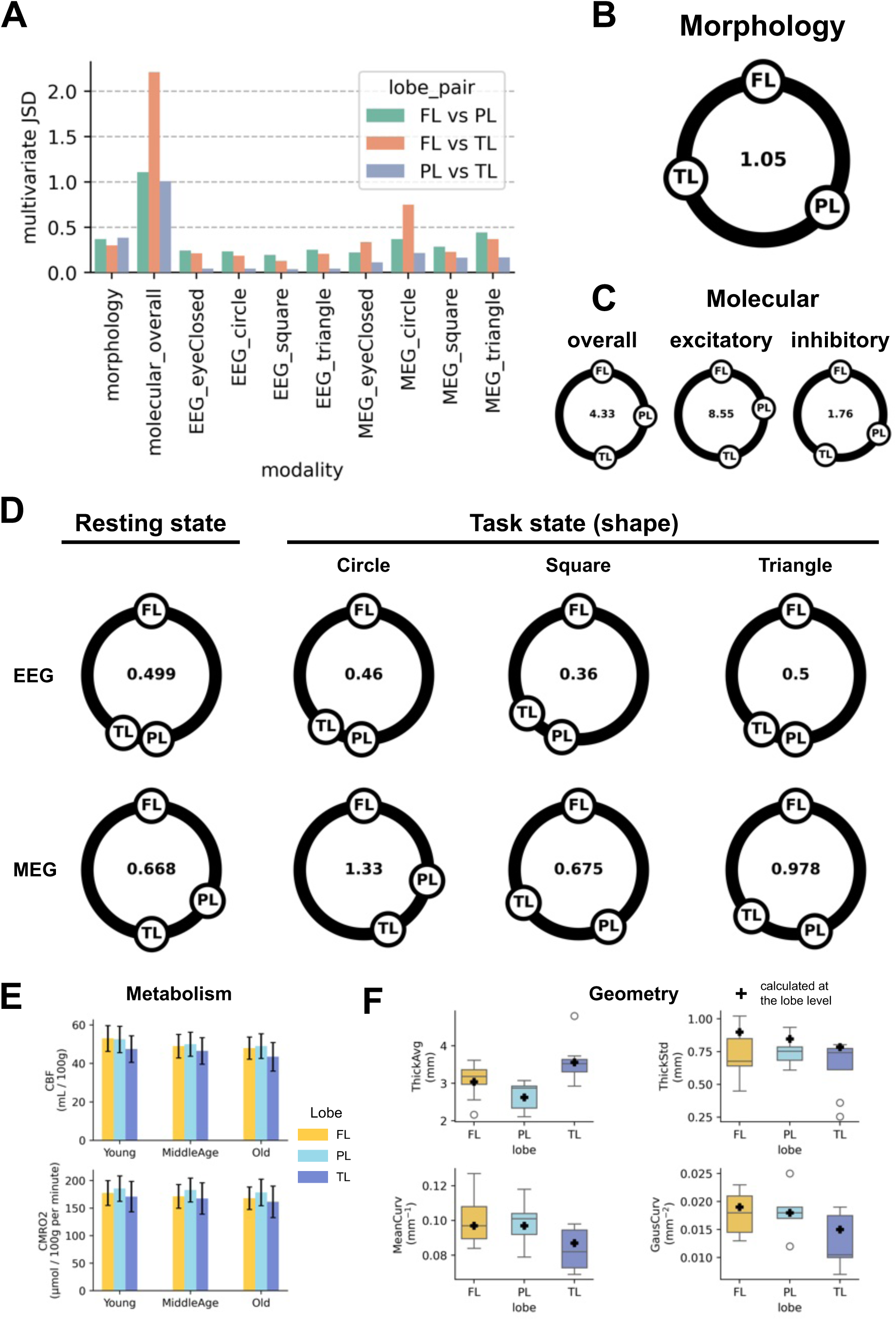
Multi-modal comparison of brain lobe separability in the human brain. **A**, Bar plot showing the MJSD (**Methods**) value for each lobe pair in different modalities (**Methods**). **B,** The relative separability between each lobe pair using neuron morphological characteristics. The length of the curve connecting two lobes indicates the relative degree of separability. Center of the circle: sum of MJSD (same convention also in **C** and **D**). **C,** Relative separability between each lobe pair based on gene expression profiles under three conditions: (1) using all neurons, (2) using only excitatory neurons (67%), and (3) using only inhibitory neurons (33%). **D,** The relative separability between each lobe pair using EEG and MEG signals. Four tasks were involved to compute the separability (**Methods**). **E,** Bar plot showing measurements of the metabolic state of brain lobes at different ages. Error bar indicates the range of mean ± standard deviation. **F,** Box plot showing measurements of the cortical geometry. The center line within each box represents the median, the lower and upper edges represent the 25th and 75th percentiles, and the whiskers represent the minimum and maximum values within 1.5 times the interquartile spacing. Outliers are shown as single hollow points. The plus symbol represents the measured value derived from the calculation at the lobe level.

## Discussion

This work provides an atlas-based framework for cross-species comparison of cortical neurons, addressing key challenges in neuron mapping, regional correspondence identification, and morphology analysis. The successful generation and alignment of large-scale datasets of mouse and human cortical neurons provide a wide coverage and quantifiable localization. By integrating anatomical, functional, and molecular information, we identified corresponding brain regions between mice and humans, facilitating a meaningful and targeted comparison of their intrinsic morphological features that has not been accomplished before.

A primary finding of our study is that human cortical neurons have significantly higher complexity compared to mouse neurons, particularly in terms of dendritic branching frequency and normalized dendritic branching intervals. The higher branching frequency and denser branching patterns of human neurons compared to mouse neurons may be directly linked to an enhanced dendritic computing capacity, a topic that warrants further investigation in future studies. These differences are pronounced in the secondary motor cortex and superior frontal gyrus, which may be related to evolutionary divergence manifested in the enhanced cognitive and motor abilities in humans over mice. While previous studies have noted differences in dendrite length and complexity on limited neurons and brain regions between species (Mohan et al., 2015; Benavides-Piccione et al.,2020; Mihaljević et al., 2021; Degl’Innocenti et al., 2023), our analyses expand on this by systematically quantifying these differences across multiple and finer cortical regions, utilizing a large-scale neuron reconstruction dataset and a carefully designed analytics approach to explore the feature space.

Our classification analysis shows that human neurons exhibit higher separability between cortical lobes compared to mouse neurons, suggesting a higher degree of morphological specialization in the human cortex. This is particularly evident in the frontal and parietal lobes, where human neurons are classified with close to 70% accuracy, compared to approximately 55% accuracy for mouse neurons. Yet, it is still unknown whether this increased morphological diversity observed in human cortical regions would reflect an elevated correlation of neuronal structural complexity with functions, which could be an interesting future topic. Moreover, local dendritic branching densities could be a feature contributing to the lobe separability, which strengthens the diversity of branching complexity in different brain regions.

From a methodological standpoint, previous efforts to annotate corresponding brain regions across species often relied on empirical methods, lacking the rigorous quantification needed for robust validation. In contrast, our analytic approach integrates uniform neuron sampling, minimal alignment error, evidence-based regional correspondence, and probabilistic neuron localization. Our approach also uses molecular profiles to validate genetic similarities between cross-species brain regions. This framework demonstrates strong generalizability, allowing for the creation of a morphological atlas of the human brain with an increasing number of neurons. Clearly, this framework may be extended to explore conserved and divergent features, not limited to neuronal morphology, in other mammalian species.

We acknowledge that there is ongoing debate about the extent to which meaningful cross-species comparisons can be made, and whether such comparisons should occur at the brain region level, the cellular level, or even the subcellular level (e.g. Reid, et al, 2016; Croxson, et al, 2018; Barron, et al, 2021; Berson, et al, 2023). An interesting future direction would involve incorporating more molecular signatures to better identify correspondences between brain regions and cell types (e.g. Berg et al., 2021; Lee, Budzillo, et al., 2021; Bakken et al., 2021; Jorstad, Song, et al., 2023; Lee, Dalley, et al., 2023). Additionally, using molecular, biochemical, and physiological approaches to determine the excitatory or inhibitory properties of cells (e.g. Siletti et al., 2023; Jorstad, Song, et al., 2023) could broaden the potential for a more accurate and comprehensive assessment of neuronal similarities and differences. Notably, exploring these additional dimensions would complement the approach presented in this work and further expand the applications of our analytical framework.

In addition to cross-species comparisons, we have examined whether the lobe separability observed in human single-neuron morphology could be reflected in other data modalities such as gene expression, EEG, MEG, cortical geometry, and metabolism. Our quantitative analysis indicates that gene expression shows the highest degree of lobe separability, highlighting the importance of molecular differences in distinguishing brain regions. In contrast, separability based on physiological signals like EEG and MEG tends to be task-dependent. However, EEG and MEG hold potential for constructing brain function models across different activity states. Metabolic features contribute less to lobe differentiation, with only subtle regional differences. Although anatomical geometry demonstrates some ability to differentiate lobes, additional brain images may be required to reach a more definitive conclusion. It is also an interesting future topic how to integrate multimodal data for classification of brain regions. Our results suggest that a comparative analysis of single-neuron intrinsic features could serve as a valuable framework for future comprehensive studies of cross-species brain structures and functions.

## Acknowledgements

This project was mainly supported by a New Cornerstone grant awarded to H.P.. The Southeast University team was also supported by a MOST (China) Brain Research Project 2022ZD0205200/2022ZD0205204 awarded to L.L.. We thank Sujun Zhao, Linus Manubens-Gil, Yufeng Liu, Feng Xiong, Shengdian Jiang for comments on the analyses and figures. We thank Ed Lein, Hongkui Zeng for discussion and contribution of some related data in this study. We also thank Javier DeFelipe, Ruth Benavides-Piccione, and Giorgio Ascoli for discussion.

## Author contributions

H.P. conceptualized and designed this study, and managed the entire project. L.Z. and N.J. performed surgeries and provided brain tissues along with metadata with the assistance of Yujin Wang, T.L., Yi Wang and Y.M.. Yujin Wang and T.L. participated in tissue collection, processing, and transportation, all clinical and imaging data collection, and sample imaging registration and matching. Z.Y. conducted all data processing and analyses under the detailed instruction of H.P.. S.D. provided initial anatomical correspondences. H.X. provided ideas for cross-species transcriptome analysis and implemented the integration of single-cell data. Y.L. designed the experiments and performed feature extraction of EEG and MEG data. B.S. and H.Z. collected EEG and MEG data. H.Z. performed data preprocessing of EEG and MEG data.

N.M. measured the anatomical properties of brain structures. R.W. performed human brain registration with the help of Z.Y.. J.R. led human cell injection, where W.Y. made a major data collection. X.C. produced human neuron reconstructions. M.D. and N.J. helped the MEG data acquisition with Y.L., B.S., H.Z.. L.L. co-managed the acquisition of human neuron morphology and reconstruction of mouse neuron morphology. L.Z. coordinated the human *ex vivo* surgical tissues and advised on the human related study. H.P. and Z.Y. wrote the manuscript with input from all coauthors.

## Methods

### Nomenclature and abbreviation of brain regions and areas in human and mouse brains

#### Human brain regions

- Temporal Lobe (TL): superior temporal gyrus (STG), middle temporal gyrus (MTG), inferior temporal gyrus (ITG), transverse temporal gyrus (TTG), entorhinal, parahippocampal gyrus (PHG), fusiform gyrus (FuG).

- Frontal Lobe (FL): superior frontal gyrus (SFG), middle frontal gyrus (MFG), inferior frontal gyrus (IFG), orbitofrontal cortex (OFC), lateral orbitofrontal cortex (OFCl), medial orbitofrontal cortex (OFCm), precentral gyrus (PrCG).
- Parietal Lobe (PL): postcentral gyrus (PoCG), supramarginal gyrus (SMG), superior parietal lobule (SPL), inferior parietal lobule (IPL), precuneus (PrCun).
- Occipital Lobe (OL): lingual gyrus (LiG), pericalcarine (PERI), cuneus (Cun), lateral occipital (LO).
- Cingulate Cortex (CC): rostral anterior cingulate cortex (rACC), caudal anterior cingulate cortex (cACC), posterior cingulate cortex (PCC).

#### Mouse brain regions

Frontal pole (FRP), somatomotor areas (MO), primary motor area (MOp), secondary motor area (MOs), somatosensory areas (SS), primary somatosensory area (SSp), supplemental somatosensory area (SSs), gustatory areas (GU), visceral area (VISC), auditory area (AUD), dorsal auditory area (AUDd), primary auditory area (AUDp), posterior auditory area (AUDpo), ventral auditory area (AUDv), postrhinal area (PoR*), anterior cingulate area (ACA), prelimbic area (PL), infralimbic area (ILA), orbital area (ORB), agranular insular area (AI), retrosplenial area (RSP), posterior parietal association areas (PTLp), temporal association areas (TEa), perirhinal area (PERI), ectorhinal area (ECT), entorhinal area (ENT).

*PoR is the correct abbreviation for Postrhinal area, which however is documented as VISpor in the Allen CCFv3. There has been debate on whether or not VISpor is correct.

#### Reconstruction of mouse local morphology

The sparsely labeled fMOST mouse brains were previously introduced by Peng et al. (2023), where the raw brain images are publicly available through the Brain Image Library (BIL, http://www.brainimagelibrary.org). The specific brain identities used in this study are listed in **Supplementary Material 1**.

The preprocessing and reconstruction followed the similar procedures described in Liu, Jiang et al. (2023). However, there were several modifications. We used 16-bit highest resolution images with a size of 1024 × 1024 × 256 voxels. We applied the NIEND (Zhao et al., 2024) image enhancement algorithm to the brain images. We only used the APP2 algorithm for neuron tracing.

#### Human specimens and reconstruction of human local morphology

A total of 23 human cases were included in this study, including 22 surgical cases from Beijing Tiantan Hospital and 1 case of donated brain tissue from the Human Brain and Tissue Bank of National Clinical Research Centre for Neurological Disorders. Informed consent was obtained in all cases, and all research procedures involving human tissues were approved by the Ethics Committee of Beijing Tiantan Hospital.

The procedures including human specimen acquisition, brain tissue preparation, sectioning, neuron labeling, neuron imaging, and neuron tracing were consistent with those documented by Han et al. (2023), but with an increase in the number of human cases and the number of neuron reconstructions. Four more human cases (H020, H021, H022, H023) acquired from surgical procedures in Beijing Tiantan Hospital were enrolled in this study. The detailed information on extracted brain tissues from each patient was summarized in **Extended Data Table 1**.

#### Mouse brain registration and neuron mapping

Brain images were aligned to 25 μm CCFv3 templates using the cross-modal alignment tool mBrainAligner (Qu et al., 2022; Li et al., 2022). We used a similar approach described in these papers. Registered channels were utilized whenever possible. Prior to alignment, the resolution of the brain was reduced to approximately 25 μm. Non-brain tissue was semi-automatically removed using Vaa3D (Peng, Ruan, Long et al., 2010; Peng et al., 2014; Liang et al., 2023). Neuronal local morphologies were inversely mapped into CCFv3 space using the inverse deformation field.

#### Human brain registration and sampling region mapping

All subjects involved in this study had complete MRI data. On the basis of the operation records and the preoperation and postoperation MRI data, experienced surgeons provided the annotated segmentation indicating the neuron sampling regions. Preference was given to using preoperative T1 MR images of the subject’s brain for alignment and sampling region outlining.

The neck was first cropped on most images and retained complete brain tissue. Then we automatically stripped skulls utilizing the Brain Surface Extractor (BSE) tool in the BrainSuite (version 23a). The results of skull stripping were refined by BrainSuite if needed. Next, the brain images were registered onto the ICBM 2009c Nonlinear Asymmetric template (a template of ICBM 152 Nonlinear atlases) using mBrainAligner. Segmentations of neuronal sampling regions were inversely mapped into ICBM 152 space using the inverse deformation field.

#### Neuron morphology standardization

The optimal neuron crop radius was set to 50 μm based on the average extension range of human neuronal Z-axis reconstruction (**Extended Data Fig. 1A**).

We calculated the number of primary branches after assembling branches within a spherical range (1 to 20 μm). Using the number of primary branches from neurons in the same brain regions in SEU-ALLEN1741 as a reference, we calculated the Jensen-Shannon Divergence (JSD) between the distribution of primary branches in the processed data and that in the reference data. Based on the smallest JSD value, we determined the optimal radius to be 8 μm for correcting the soma topology. (**Extended Data Fig. 1B**)

The reconstructions in SWC format were resampled to 5μm spacing between continuous nodes.

#### Morphological features

The following morphological features are defined in L-Measure (Scorcioni et al., 2008):

- ’Avg. Straightness’ (known as ‘Avg. contraction’ in L-Measure’s nomenclature), ‘Avg. Remote Bifur Angle’, ‘Max Branch Order’, ‘Bifur Num’ (bifurcation number), ‘Total Length’, ‘Max Path Dist’, ‘Avg. Euclidean Dist’, ‘Avg. Path Dist’.
- ‘Center Shift’ is defined as the distance between soma and the centroid of the neuron which is computed by the weighted average of the spatial position of all nodes in neuron reconstruction. Nodes are weighted by the distance to its parent node. Equal weight is assigned to each node if a neuron reconstruction is resampled with a fixed distance of conterminous nodes.
- ’Volume’ is defined as the volume of a convex hull in a three-dimensional space. The convex hull is calculated by the ‘scipy.spatial.ConvexHull’ function (SciPy, Python package, version 1.9.1). Our measuring unit is μm^3^. “Volume” was log_10_ transformed in analysis.
- ’3D Density’ is defined as the ratio of ‘Total Length’ and ‘Volume’. “3D Density” was log_10_ transformed in analysis.
- ’BO[X] Branch Num’ denotes the number of branches with the Xth branch order, X=1∼5.
- ’BO[X] Branch Length’ denotes the sum of lengths of branches with the Xth branch order, X=1∼5.

#### Acquisition of RNA-seq data from mouse and human brain

Single-cell RNA-seq data of the mouse brain were generated by Yao et al. (2023). These data are available at the Neuroscience Multi-omics Archive under identifier: https://assets.nemoarchive.org/dat-qg7n1b0.

Single-nucleus RNA-seq data of the human brain were generated by Siletti et al. (2023). These data are available at the CELLxGENE: https://cellxgene.cziscience.com/collections/283d65eb-dd53-496d-adb7-7570c7caa443.

#### Data filter and quality control

Human snRNA-seq data were filtered by three criteria: a. retained the cells sampled from cortical regions provided in the metadata of human gene data. The region contains A1C, A23, A40, A44-A45, A46, A5-A7, A24, ITG, M1C, MTG, Pro, S1C, STG, TF, TH-TL, V1C, V2.

According to Table S4 of the “Supplementary Materials” section in this work (Siletti et al., 2023), these dissection names corresponded to the following anatomical regions: TTG, PCC, SMG, IFG, MFG, SPL, rACC, ITG, PrCG, MTG, Cun, PoCG, STG, FuG, PHG, Cun-LiG, Cun. b. according to the “supercluster_term” column in the cell metadata, we retained the cells in the following classes: “Upper-layer intratelencephalic”, “Deep-layer intratelencephalic”, “MGE interneuron”, “CGE interneuron”, “Deep-layer corticothalamic and 6b”, “LAMP5-LHX6 and Chandelier”, “Deep-layer near-projecting”, “Miscellaneous”. c. retained the cells labeled “Y” in column “Sample retained in analysis” in the cell metadata.

Mouse scRNA-seq data were filtered by two criteria: a. retained the cells sampled from cortical regions named “Isocortex” in the cell metadata. b. removed cells clustered into non-neuronal classes. Non-neuronal classes contain “Astro-Epen”, “OPC-Oligo”, “OEC”, “Vascular”, “Immune”, “DG-IMN Glut”, “OB-IMN GABA”, “Pineal Glut”, “CB Glut”.

Both human and mouse genetic data removed cells with a mitochondrial gene percentage greater than 5%. Finally, 299,219 mouse cells and 648,730 human cells were involved in analyses.

#### Data integration

The overall integration was conducted by Seurat v5 (R package, version 5.0.3) (Hao et al., 2024), following the procedure described below: Gene counts of each cell were normalized to a total of 10,000 and then applied natural logarithm. The process was implemented using the NormalizeData function with default parameters. The top 2,000 highly variable genes (HVGs) were identified using the FindVariableFeatures function with default parameters. To manage computational load, we sketched the merged dataset to 5,000 cells using the LeverageScore method implemented in the SketchData function. The sketched data were integrated using Canonical Correlation Analysis (CCA) through the IntegrateLayers function. Finally, dimensional reductions learned on the sketched cells were extended to the full dataset.

#### Cell-type-count coding

We calculated K nearest neighbors (n_neighbors=20, n_pcs=10) using a dimensional reduction of Seurat CCA. Leiden clustering (Traag et al., 2019) was applied to the integrated full dataset using SCANPY (Python package, version 1.9.8) (Wolf et al., 2018) and generated 25 clusters. We eliminated clusters that had less than one-thousandth of the total number of cells (threshold: 299 for mice and 648 for humans). Then cells from each brain region were counted into 21 clusters to get a cell-type-count coding.

#### Correlation of genetic profiles

The sum of each cell-type-count coding was normalized to 1 and then z-score standardized by cluster within the respective species. Principal component analysis (PCA) was then utilized to obtain denoised representations with the number of principal components ranging from 7 to 14 (corresponding to variance explained ratios of 0.90 to 0.99, respectively). Pairwise similarities and deviations could be calculated by the average and standard deviation of the Pearson correlation coefficient between 8 PCA representations. The confidence was defined as the maximum of deviations subtracting each deviation. PCA (function “decomposition.PCA” in scikit-learn Python package, version 1.2.2). Pearson correlation coefficient (function “stats.pearsonr” in SciPy Python package, version 1.9.1)

#### Integrated ranking of genetic similarity

Besides the average Pearson correlation coefficient described in the last section, we also calculated the cosine similarity based on each PCA representation. Additionally, according to the cell-type-count coding of each brain region, we constructed the cell-type-count matrices for each species. We used chi-square tests to calculate the cell types enriched in each brain region. We calculated the ratio of the number of observed cells to the expected number of cells (R*_o/f_*) for each cluster in each species (Cheng et al., 2021). We recorded the first M cell types with the highest R*_o/f_*) values in each brain region and calculated the Jaccard index between different brain regions. We sorted all three similarity metrics (correlation coefficient, cosine similarity, Jaccard index) in descending order respectively, and recorded the average rankings between a particular region and every other region of another species.

#### Weighted statistical measures

We assigned each neuron’s probability of belonging to each region (**Fig. 3**, **Extended Data Fig. 3A**). The sum of the probabilities of all cells in each region was then normalized to 1.

Let *X* be the matrix of morphological features, where *X_ij_* denotes the value of the *j*-th feature for the *i*-th neuron. Let *W* be the matrix of soft labels, where *W_ij_* represents the probability of the *i*-th neuron being assigned to the *j*-th brain region. The Weighted Mean for the *j*-th feature was computed as follows:

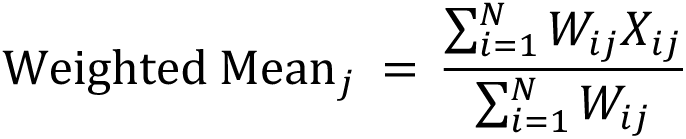

where *N* is the total number of neurons.

For each feature *j*, the values *X_ij_* for all neurons *i* were sorted in ascending order, resulting in a sorted list of feature values *X_j_*. The corresponding weights *W_ij_* were also sorted to match the sorted feature values. The cumulative weights were calculated, and the Weighted Median was identified as the feature value corresponding to the point where the cumulative weight reached 0.5 (**Extended Data Fig. 3B**).

Similarly, the first weighted quartile (Q1) and third weighted quartile (Q3) were computed by identifying the feature values at the cumulative weights of 0.25 and 0.75, respectively.

#### Weighted statistical test

We performed statistical comparisons between groups using a weighted Kolmogorov-Smirnov (KS) test. The analysis was conducted in the R environment utilizing Ecume package (version 0.9.2) for each test (Roux de Bézieux et al., 2024).

#### Quantification of feature separability

The features sorted by mRMR (pymrmr, Python package, version 0.1.11) are sequentially used in classification. We standardized the morphological features using z-score normalization. Dimensionality reduction was then performed using PCA, selecting principal components that collectively explained 95% of the variance. We employed a 5-fold cross-validation technique and made folds retain the percentage of samples in each category when splitting the dataset. For classification, we utilized a Random Forest classifier (ensemble.RandomForestClassifier function in scikit-learn Python package, version 1.2.2) with fixed parameters: max_features = “sqrt”, class_weight = “balanced”. A grid search was conducted to optimize the hyperparameters, specifically examining the following ranges: n_estimators = [100, 200, 300, 400, 500], min_samples_split = [2, 3, 4], and max_depth = [4, 5]. The model’s performance was evaluated using the balanced accuracy score, and the configuration yielding the highest average test score was selected as the optimal model.

#### Multivariate Jensen–Shannon divergence (MJSD)

For two multivariate distributions P_1_ and P_2_ with mean vectors μ_1_, μ_2_ and covariance matrices Ʃ_1_, Ʃ_2_, the JS divergence is computed as:

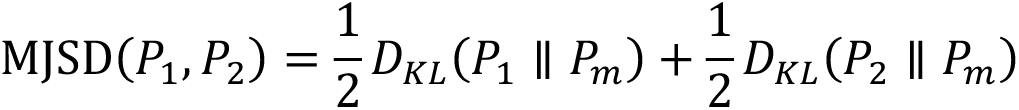

Where *P_m_* is the multivariate distribution with mean vector μ_m_ and covariance matrix Ʃ*m*::

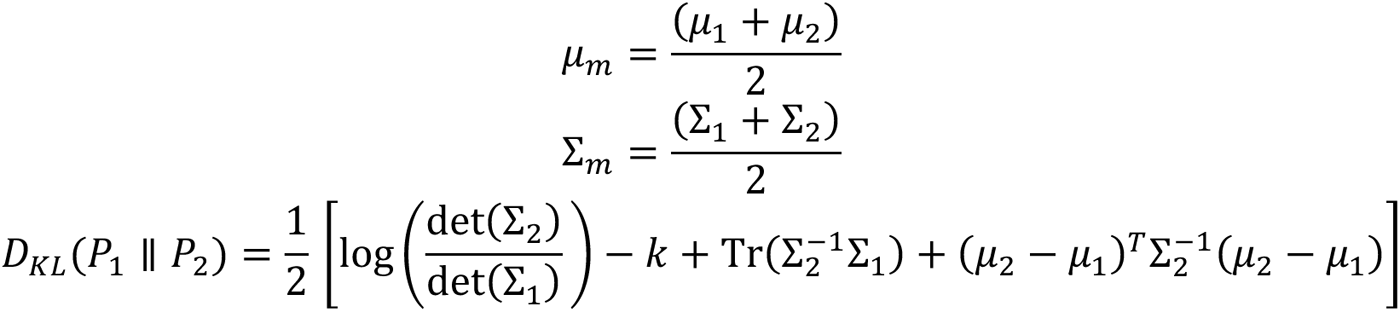

### Multimodal lobe separability

#### Cortical geometry

We used FreeSurfer (version 7.4.1; Fischl, 2012) software to measure the anatomical geometry of brain structures by running the “recon-all” command. The region-level statistics were derived from the “rh.aparc.stats” file. We also ran the “mri_annotation2label” command with the “--lobesStrict” parameter to get a lobe annotation. Then we re-called the “mris_anatomical_stats” command to generate the geometric statistics at the lobe level. We verified the structural segmentation using Freeview, a visualization tool in FreeSurfer.

#### Physiological signals

**Data acquisition:** We recorded synchronized EEG and MEG data (collectively known as MEEG) from 9 subjects, aged 22 to 28, during resting and task states. The group comprised 3 females and 6 males. Each subject was placed in a dark room and equipped with a 61-sensor MEG recording helmet and a 22-sensor EEG system. The MEG sensors were mounted on a helmet fixed to an operation bed in a magnetically shielded environment, where the subjects were instructed to lie down. The EEG sensors were applied using the 10-20 system under medical supervision, with an additional sensor placed on the chest to monitor the electrocardiogram (ECG). The ground electrode was attached to the left arm, while the reference electrode was placed on the prefrontal cortex. Both EEG and MEG signals were sampled at a rate of 1000 Hz. Initially, subjects were asked to close their eyes, relax, and maintain a resting state for approximately 2 minutes while the MEEG data were recorded. Each 2.8-second interval was treated as a separate trial. In the task state, participants were instructed to perceive and imagine a random shape, with each trial consisting of 1-second of preparation, 0.2-second of perception, and 1.6-second of post-imagery. The sizes of shapes were equal. There were 540 trials for each shape.

**Data preprocessing:** The MEEG data were preprocessed using the MNE-Python package (version 1.7.1). We automatically processed all trials, starting with filtering the data using a passband of 2–80 Hz, applying a 50 Hz notch filter for EEG data, and a 22 Hz notch filter for MEG data. Baseline correction was performed based on the preparation period, and channels with relative amplitudes exceeding 0.0004 V for EEG and 40 fT for MEG were excluded. Next, the data were decomposed into 15 independent components using independent component analysis (ICA), and artifact components representing heartbeats and eye blinks were manually identified and removed.

**Feature extraction:** For each trial, we extracted 13 features used for the calculation of MJSD from both EEG and MEG signals. These features included latency of maximum value, latency of minimum value, latency of maximum amplitude, standard deviation, skewness, kurtosis, differential entropy, zero-crossing ratio, outlier ratio, and four band power ratios. The power ratios corresponded to the following frequency bands: delta (0.1–4 Hz), theta (4–8 Hz), alpha (8–14 Hz), beta (14–30 Hz), and gamma (30–100 Hz).

**MEEG sensor labeling:** The 21 EEG sensors are of the international 10-20 sensor placement system and the 61 MEG sensors were designed by Beijing X-MAGTECH Technologies. All the sensor coordinates were registered to ICBM 2009a Nonlinear Symmetric 1×1×1 mm template approximately. Then we performed edge detection on the brain mask of the ICBM template and identified a total of 198128 edge voxels as the sources. To evaluate the electric or magnetic field contribution, we calculated the Euclidean distance between each sensor coordinate and each voxel. We selected the nearest 100 voxels, and the CerebrA label of the brain region with most enrichment was noted as the label of one sensor.

#### Metabolism

Data on global metabolic status were taken from Table 2 of Aanerud et al. (2012). Both mean and standard deviation were averaged by the left and right brain hemispheres.

#### Molecular

Molecular data were collected using human snRNA-seq data (Siletti et al., 2023). The cells in the frontal, parietal, and temporal lobes were downsampled to 10% of the original data according to their superclusters labeled in the dataset. The genes on the X and Y chromosomes were excluded. A series of genes associated with mitochondria, heat-shock proteins, ribosomes, and dissociation were also excluded. The latter gene list was taken from Supplementary Table 1 of Xue et al. (2022). Excitatory neurons were classified under the following categories in the “supercluster_term” column: “Upper-layer intratelencephalic”, “Deep-layer intratelencephalic”, “Deep-layer corticothalamic and 6b”, and “Deep-layer near-projecting”. Inhibitory neurons were categorized as “MGE interneuron”, “CGE interneuron”, and “LAMP5-LHX6 and Chandelier” in the same column. We processed the data using the SCANPY (version 1.9.8) Python package. The gene count matrix was sequentially applied with normalize_total, log1p, highly_variable_genes (5,000 HVGs), and pca (50 PCs) functions. The top 50 PCs were used to calculate the MJSD.

## Data and materials availability

The mouse neuron reconstructions can be accessible on Zenodo (https://doi.org/10.5281/zenodo.13801372).

The human neuron reconstructions and other Supplementary Materials can be accessible on Zenodo (https://doi.org/10.5281/zenodo.13801283).

## Supplementary Materials

**Extended Data Fig. 1.**
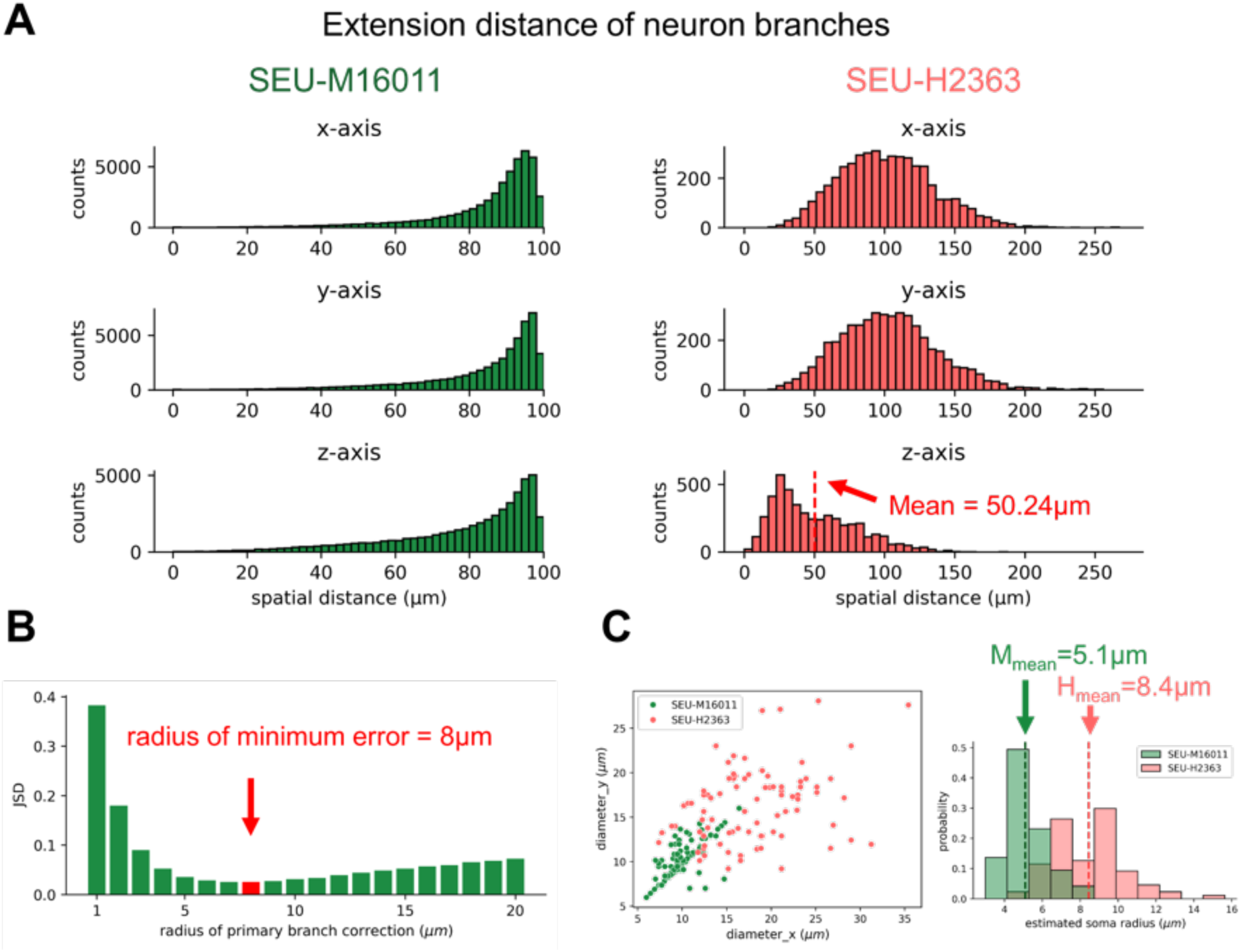
Basic statistical information on neuron reconstructions. **A**, Histograms of neuronal maximum tracing distance from soma along three axes for mouse (SEU-M16011) and human (SEU-H2363) neuron reconstructions. The mean distance is labeled for human data on the z-axis chart. **B,** Bar plot showing JSD of primary branch numbers between SEUALLEN1741 cortical neurons and SEU-M16011 neurons under different assembling radii. The bar of the minimum JSD is labeled in red (x = 8μm). C, Statistics of soma size. Scatter plot: soma diameters of 100 randomly selected neurons along the x-axis and y-axis. Histogram: distribution of estimated soma radius of human and mouse data.

**Extended Data Fig. 2.**
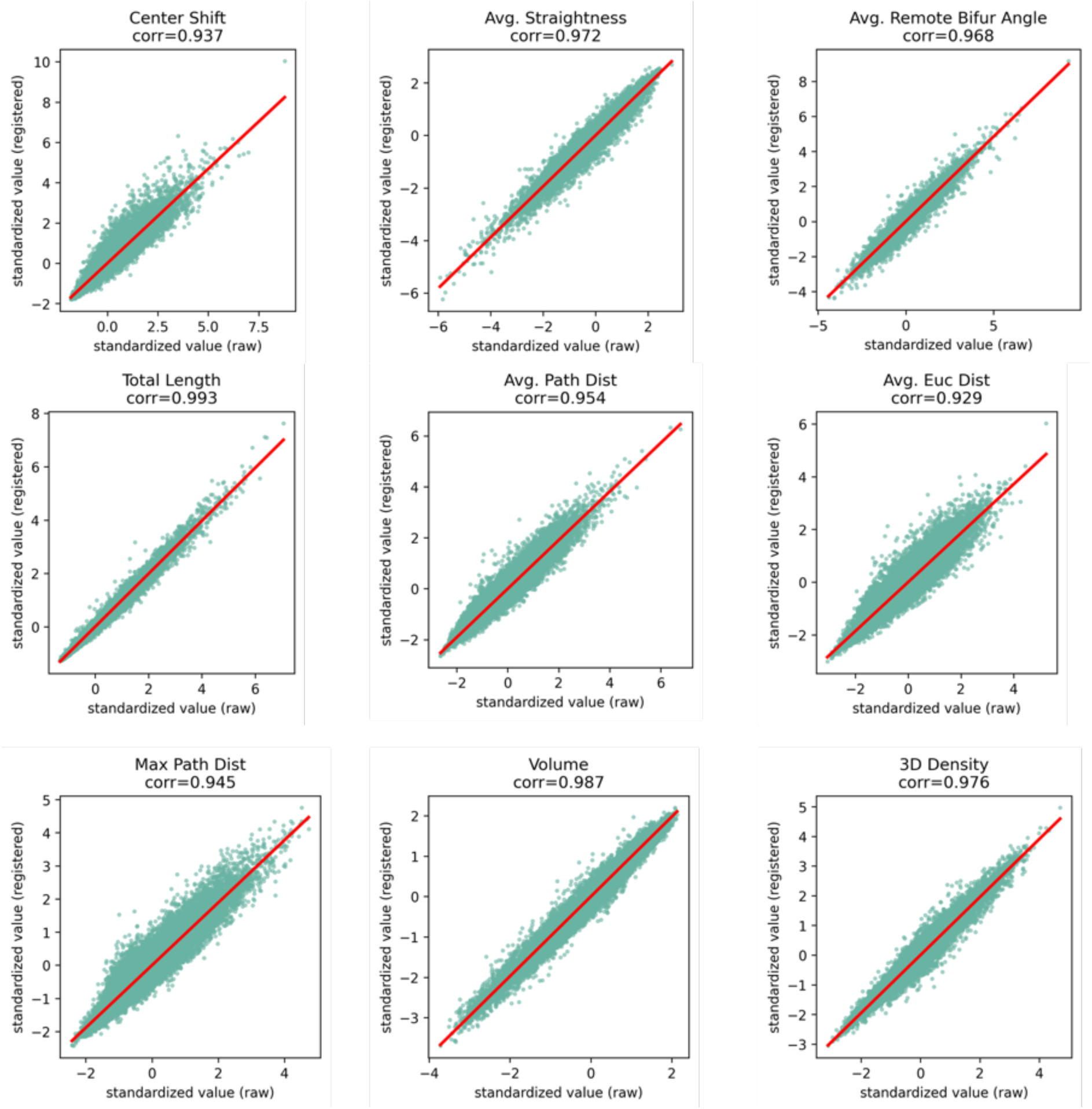
Difference in morphological features between non-registered (raw) and registered neuronal reconstructions. X-axis: feature value by z-score standardization of non-registered neurons. Y-axis: feature value by z-score standardization of registered neurons. The red line denotes the linear regression fitted result. The Pearson correlation coefficient is labeled below the feature name.

**Extended Data Fig. 3.**
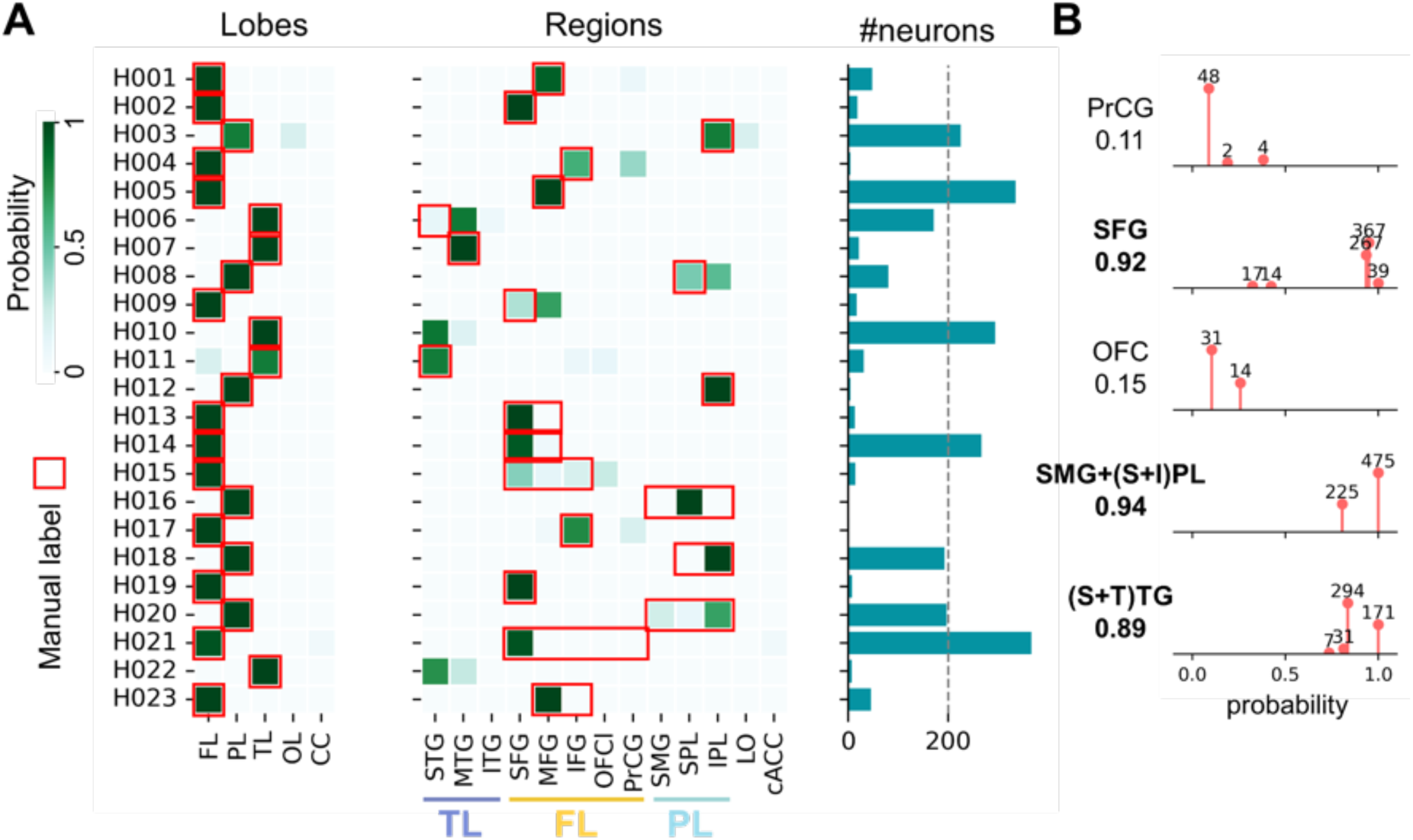
Summary of the alignment results for human sampling tissues. **A**, Statistics of the regional proportions of individual sampling tissues. Left heatmap: proportion of individual sampling tissues at the lobe level. Middle heatmap: proportion of individual sampling tissues at the level of finer anatomical regions. Color bar: proportion of tissue samples belonging to a brain region after alignment. The red boxes represent the expert’s annotations. Cases of H010 and H022 are not labeled in the red box because they were manually labeled as temporal poles; however, the Mindboggle-101 atlas has reassigned the temporal pole to the superior temporal gyrus, middle temporal gyrus, and inferior temporal gyrus. Right bar plot: the total number of neurons from each subject. The reason for the H006 mismatch is the low imaging resolution. The resolution of the MRI image is 512 × 512 × 23 voxels, where the first two dimensions correspond to the axial planar resolution and the third dimension represents 23 slices acquired along the dorsal-ventral axis. This case was manually labeled as superior temporal gyrus with probability 1. **B,** Lollipop chart showing the composition of neurons in brain regions with partially established correspondences. X-axis: probability of neurons belonging to the region(s). The height of lollipops denotes the number of neurons. The region name and average belonging probability are labeled on the left, corresponding to the human brain region in Fig. 3E. Selected regions for subsequent analyses are marked in bold font. Only regions with neurons are listed.

**Extended Data Fig. 4.**
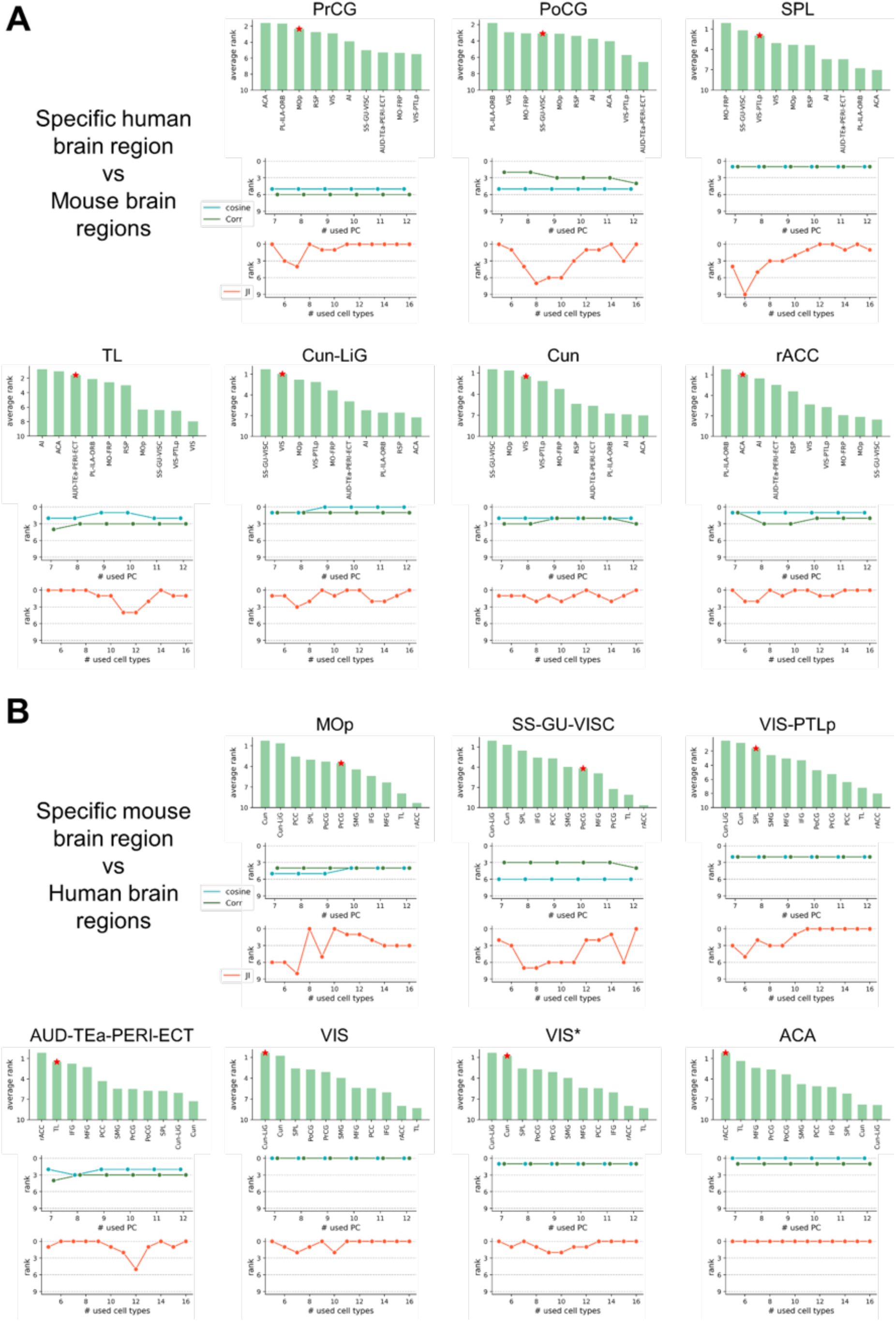
Genetic correspondence between mouse and human brain regions. **A**, Similarity of cell-type composition between the specific human brain region and all mouse brain regions. Bar plot: average ranks (**Methods**) between the specific human brain region and all mouse brain regions, sorted by ascending order. The red star denotes the best match of the functional and anatomical corresponding region. Line graph: Ranking of similarity between human brain regions and their corresponding rat brain regions in terms of similarity between human and all rat brain regions under different similarity metrics. The more similar the higher the ranking. Cosine: cosine similarity; Corr: Pearson correlation coefficient; JI: Jaccard index. **B,** Similar to that in **Panel A**, but focusing on the similarities between the specific mouse brain region and all human brain regions. The second VIS region is marked with an asterisk, which indicates a one-to-many relationship (VIS and Cun-LiG; VIS and Cun).

**Extended Data Fig. 5.**
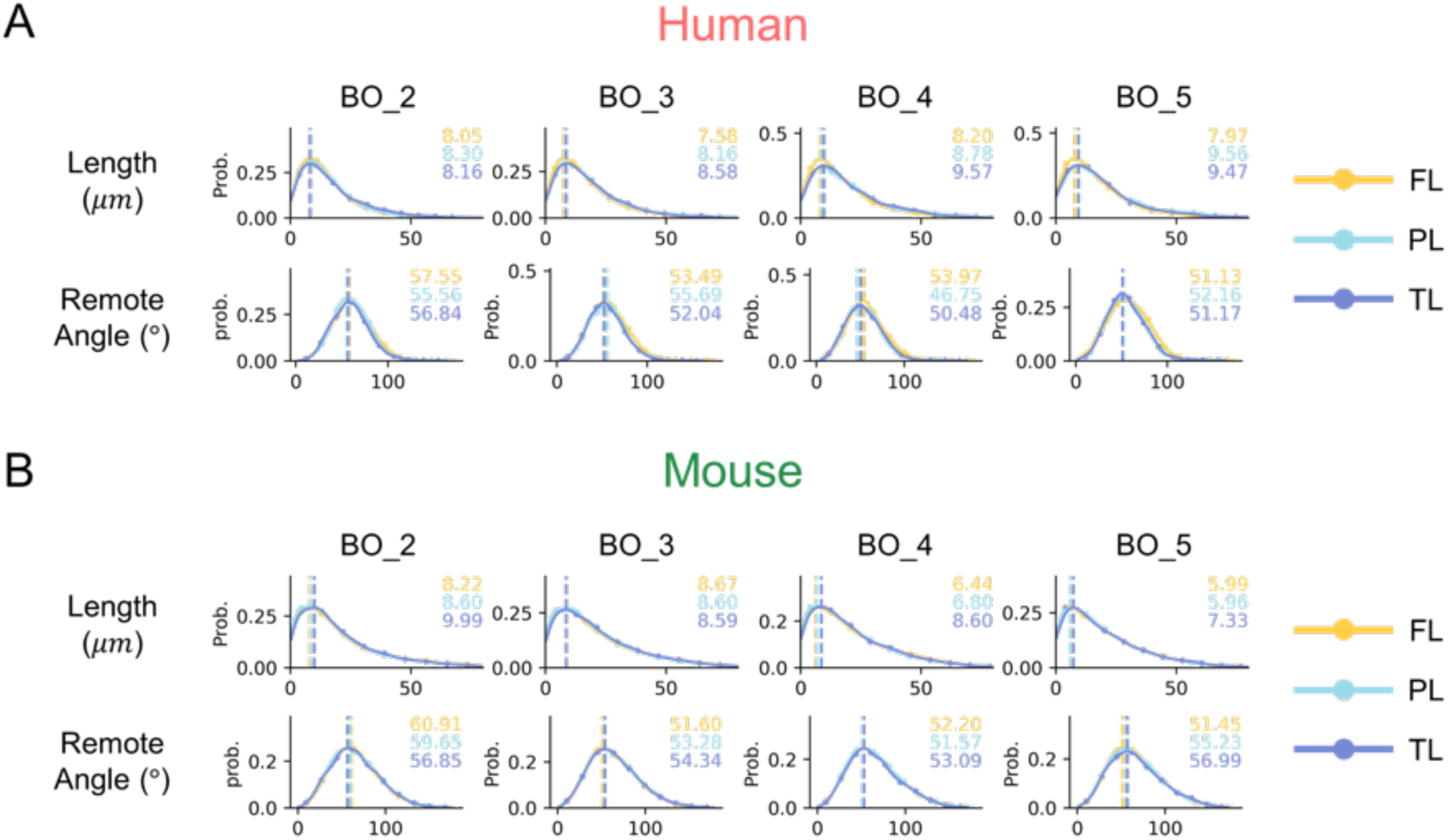
Distribution of non-terminal branching features of neurons in different brain lobes within species. **A**, Feature distribution of human neuronal branches with branch order 2 to 5 in the frontal, parietal, and temporal lobes. The dots represent the height of each bin in the histogram. The line is fitted by Gaussian kernel density estimation. The branch length and the remote angles between sibling branches were examined. The peak of each distribution is marked by a dashed line and labeled in the upper right corner. **B,** Same content as in **Panel A**, but change the species to the mouse.

**Extended Data Fig. 6.**
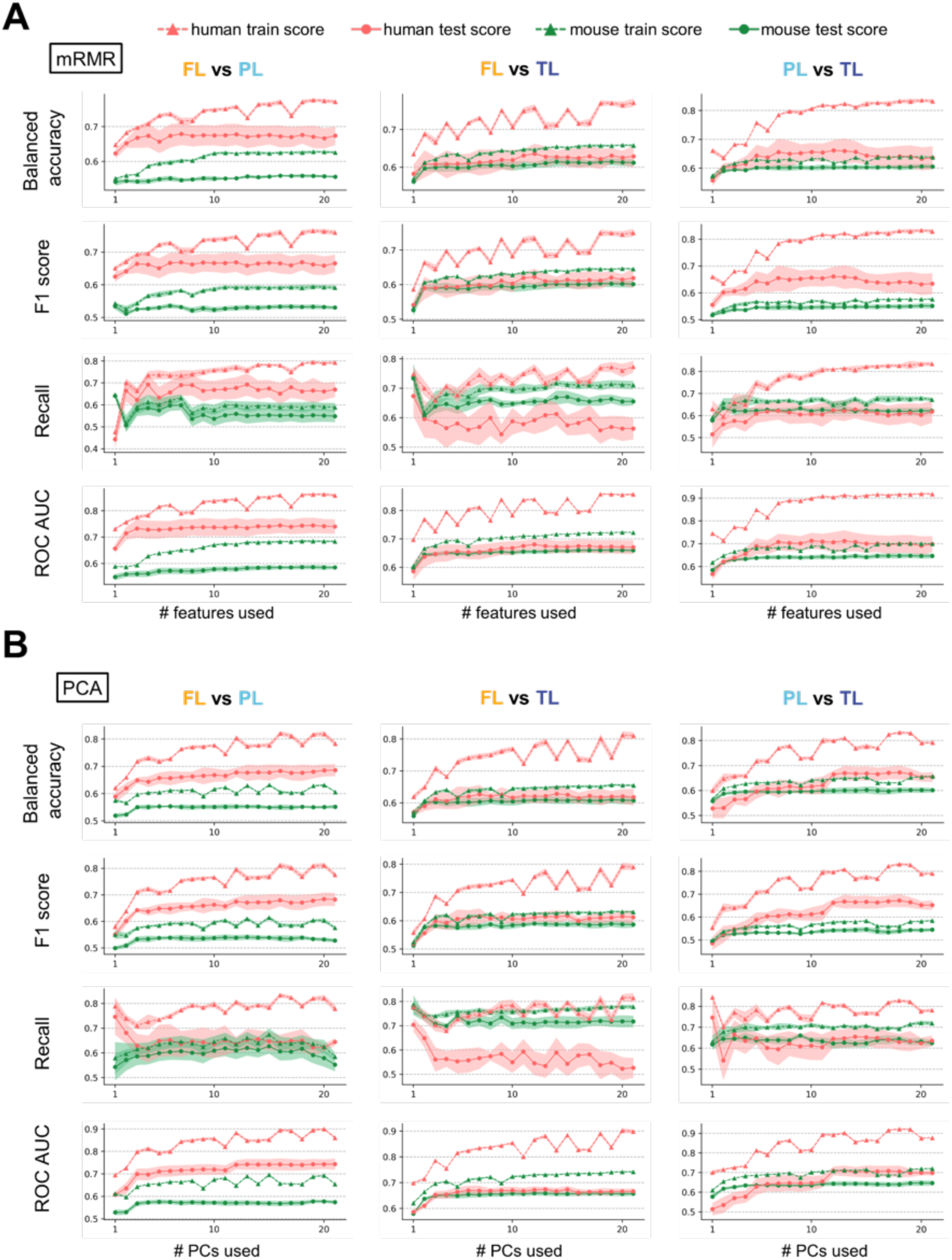
Performance evaluation of classification of neuronal morphology across different brain lobes. **A**, Classification Performance Evaluation with mRMR-based feature selection. x-axis: number of incrementally used morphological features selected by the mRMR algorithm. y-axis: the value of evaluating metric. The evaluation of both the training data and the testing data in the five-fold cross-validation is shown, with the shaded area representing the range of mean ± standard deviation. Four Metrics were involved including balanced accuracy, F1 score, recall, and area under the receiver operating characteristic curve (ROC AUC). **B,** Classification Performance Evaluation with PCA-based feature selection. x-axis: number of incrementally used Principal Components (PCs) derived from PCA applied to the complete set of morphological features. Other elements are the same as in **panel A**.

